# Vesicle-mediated mitochondrial clearance underlies an actionable metabolic vulnerability in triple-negative breast cancer

**DOI:** 10.1101/2024.04.04.587899

**Authors:** Jody Vykoukal, Yihui Chen, Monica J. Hong, Mingxin Zuo, Hansini Krishna, Hiroyuki Katayama, Ehsan Irajizad, Ranran Wu, Ricardo A. León-Letelier, Jennifer B. Dennison, Angelica M. Gutierrez, Adriana Paulucci-Holthauzen, Timothy C. Thompson, Leona Rusling, Yining Cai, Fu Chung Hsiao, Soyoung Park, Banu Arun, Samir Hanash, Johannes F. Fahrmann

## Abstract

Selective autophagy of mitochondria is known to promote survival and progression of cancer cells in various malignancies including triple-negative breast cancer (TNBC). Here, we aimed to identify the essential metabolic adaptations that support mitochondrial quality control with the goal to uncover actionable metabolic vulnerabilities with therapeutic potential. Using an integrated approach of proteomics and untargeted and stable-isotope resolved metabolomics, coupled with functional experimental analyses, we define an alternative mechanism to mitophagy enabled by an onco-metabolic program of heightened extracellular sphingomyelin salvaging in TNBC that facilitates extracellular vesicle (EV)-mediated intracellular clearance of mitochondrial damage. Targeting of the cancer cell sphingolipid onco-metabolic pathway via repurposing of eliglustat, a selective small molecule inhibitor of glucosylceramide synthase (UGCG), resulted in ceramide-induced lethal mitophagy and attenuated tumor growth and prolonged overall survival at clinically achievable doses in an orthotopic syngeneic mouse model of TNBC. Our study defines a mechanism of aberrant sphingolipid metabolism that underlies an actionable metabolic vulnerability for anti-cancer treatment.

## INTRODUCTION

Mitochondria are semi-autonomous intracellular organelles that are highly dynamic and responsive to metabolic and extracellular insults, undergoing continuous changes in their number and morphology to maintain mitochondrial homeostasis.^1^ Mitochondria play central roles in cancer development and progression by contributing to cell proliferation, apoptosis, calcium ion storage, energetics, and production of reactive oxygen species (ROS).^1^ Mitochondrial dynamics in cancer cells are regulated through a complex network of proteins that promote fission and fusion events as well as mitophagy, which plays a pivotal role in clearance of damaged/dysfunctional mitochondria to prevent excessive buildup of toxic products that may otherwise elicit deleterious effects.^1^ Removal of damaged mitochondria through mitophagy requires both induction of general autophagy and priming of damaged mitochondria for selective autophagic recognition.^2^ General induction of autophagy is mediated through autophagy-related genes (ATGs); whereas mitochondrial priming is mediated either through the Pink1-Parkin signaling pathway or through the mitophagic receptors Nix and Bnip3.^3,4^ Recent studies have also demonstrated that removal of damaged mitochondria can undergo Parkin-dependent sequestration into Rab5-positive early endosomes via the endosomal sorting complexes required for transport (ESCRT) machinery and delivered to lysosomes for degradation.^5^ This process involves maturation of early endosomes into late endosomes, via switching from Rab5 to Rab7 ^6^, prior to fusing with the lysosome for degradation.^5^ An alternative mechanism of mitochondrial quality control involves the generation of mitochondria-derived vesicles (MDVs). MDVs are generated through selective incorporation of mitochondrial protein cargoes, including oxidized proteins ^7,8^, and MDVs can be either targeted to the late endosome/lysosome for degradation via Pink1/Parkin-dependent mechanisms ^9^ or to the peroxisomes.^10^ More recent data in cardiovascular disease has demonstrated that inhibition of mitophagy promotes secretion of extracellular vesicles (EVs) containing intact mitochondria as an alternative mechanism for mitochondrial clearance.^11^ While these studies provide insights into the molecular machinery ascribed to mitochondrial quality control and turnover, the metabolic underpinnings that enable mitochondrial degradation and clearance remain poorly understood. Elucidating metabolic adaptation(s) in cancer cells that enable mitochondrial quality control may expose metabolic vulnerabilities that are targetable for anti-cancer treatment.

Triple-negative breast cancer (TNBC) is characterized by high pathological grade, strong invasiveness, local recurrence, high metastasis rate, and poor prognosis.^12^ Compared to other subtypes of breast cancer, mitochondria in TNBC exhibit a more fragmented morphology, which often precedes mitophagy ^13,14^. Protein expression of microtubule-associated protein 1 light chain 3B (LC3B), which is involved in autophagosome formation and a marker of autophagy, has been associated with disease progression and poor disease-free survival and worse overall survival in patients with TNBC.^15^ Some studies suggest that TNBC progression may be linked to defective mitophagy rather than induction of mitophagy. Specifically, studies using the MMTV-PyMT TNBC mouse model demonstrated that loss of Bnip3 is associated with TNBC development. Mechanistically, reduced Bnip3 expression in tumors resulted in defective mitophagy with consequent increases in mitochondrial mass and ROS that activated HIF-1α and downstream target genes to promote tumor development.^16^ The Bnip3 locus at chromosome 10q26.3 is also frequently deleted in metastatic progression of TNBC.^17^ Yet, Bnip3 expression is retained in early-stage TNBC ^16^ and targeting of autophagy/mitophagy by small molecules such as chloroquine and isorhamnetin in TNBC cells promotes apoptosis ^18^, suggesting mitophagic flux is active in these cells during early stages of disease. Moreover, a recent immunohistochemical analysis of autophagy/mitophagy markers in a breast cancer tissue microarray demonstrated elevated protein expression of Beclin-1, LC-3, BNIP-3, and Parkin in breast cancer tumors compared to adjacent control tissues. Notably, in this study, tumor staining positivity for BNIP3, LC3, and Beclin-1 was highest among hormone receptor (HR) negative breast cancers.^19^

In this study, we aimed to elucidate the underlying metabolic networks that enable clearance of damaged mitochondria using TNBC as a model system. Using an integrated systems approach consisting of in-depth proteomics, untargeted and stable-isotope resolved metabolomics as well as a of novel lipid particles trafficking tracking system, we detail a mechanism of mitochondrial maintenance ed an alternative mechanism alternate to mitophagy, whereby TNBC cells heighten extracellular sphingomyelin scavenging and salvaging that, in turn, facilitates EV-mediated intracellular clearance of mitochondrial damage. We further demonstrate therapeutic potential of targeting sphingolipid onco-metabolism via repurposing eliglustat, a selective small molecule inhibitor of glucosylceramide synthase (UGCG), for anti-cancer treatment in vitro and in an orthotopic syngeneic mouse model of TNBC.

## Methods

### Chemicals

Eliglustat tartate (Abmole Bioscience, M5610). Carbonyl cyanide-m-chlorophenyl hydrazone (CCCP) and carbonyl cyanide-p-trifluoromethoxyphenylhydrazone (FCCP) (Cayman Chemical, 25458 and 15218). Sphingomyelin(18:1/18:1-d9) and C11 TopFluor Sphingomyelin (N-[11-(dipyrrometheneboron difluoride)undecanoyl]-D-erythro-sphingosylphosphorylcholine) (Avanti Polar Lipids 791649 and 810265P). Azithromycin (MedChemExpress, HY-17506), Chloroquine (MedChemExpress, HY-17589A), and Bafilomycin A1 (MedChemExpress, HY-100558).

### Preparation of synthetic self-assembled lipid particles

Purified lipids were dissolved in chloroform (Avanti Polar Lipids 441601, 860584, 791649 and 700000; Sigma C9253 and T7140) and combined in ratios consistent with composition of VLDL, LDL or HDL, standardized to 6.5 mg mL^-1^ total lipids. Solvent was removed under vacuum for 18 hrs (Thermo Savant SpeedVac SC250EXP) and lipids hydrated for 30 min at 40 °C with intermittent sonication in 0.01 M HEPES / 0.15 M NaCl buffer, 1 mL per 65 mg total lipids; transferred to polycarbonate tubes and subjected to five cycles of sonication, freeze (isopropanol/dry ice) and thaw (60 °C). The resulting vesicle suspension was extruded for 11 cycles through a 100 nm nanopore membrane (NanoSizer Extruder TT-002-0010, T&T Scientific Corporation, Knoxville TN) maintained at 40 °C. Lipid particle suspensions were 0.22 µm sterile filtered and stored at −4 °C. Particles were used at 1:10–1:100 dilution. Particle size and yield was confirmed using nanoparticle tracking (ZetaView, Particle Metrix GmbH).

### Cell Lines

A panel of 34 breast cancer cell lines (AU565, HCC1419, HCC1954, HCC202, MDAMB361, SKBR3, ZR7530, MDAMB453, BT474, CAMA1, HCC1428, HCC1500, MCF7, T47D, ZR751-1, BPLER2, BPLER3, BT20, BT20, BT549, DU4475, HCC1143, HCC1187, HCC1395, HCC1806, HCC38, HCC70, HMLER3, Hs578T, MDAMB157, MDAMB231, MDAMB436, MDAMB468, and MFM223) were analysed by untargeted metabolomic profiling. All breast cancer cell lines were maintained in DMEM (Gibco, 11995065) supplemented with 10% FBS. The identity of each cell line was confirmed by DNA fingerprinting via short tandem repeats at the time of mRNA and total protein lysate preparation using the PowerPlex 1.2 kit (Promega).

### In vitro viability assays

Cell viability for TNBC cell lines following drug treatment was determined using colorimetric cell proliferation assays (MTS, BioVision Inc) per manufacturer’s instructions.

#### Confocal Microscopy

Mitochondria were imaged using BacMam CellLight^TM^ Mitochondria-RFP (fused construct of E1 alpha pyruvate dehydrogenase leader sequence and TagRFP) (ThermoFischer, C10601) and anti-TOM20-GFP antibodies (Proteintech, #CL48811802). Endosomes, and autophagosomes/lysosomes were imaged using BacMam CellLight^TM^ Late Endosome-GFP (fused construct of Rab7a), and Lysosome-GFP (fused construct of lysosomal associated membrane protein 1 and TagGFP) probes (ThermoFischer, C10588, and C10596). Mitochondrial potential was assessed using MitoTracker Red CMXRos (Cell Signaling, 9082). For immunofluorescent staining of Tom20 and Ceramide, TNBC cells treated with vehicle or eliglustat (64 µM) were fixed in 4% PFA, permeabilized in 0.1% Triton X-100, and blocked in 1% BSA, followed by overnight staining of α-Tom20 (Clone 1D6F5, Proteintech) and α-Ceramide (Clone MID 15B4, Sigma-Aldrich) antibodies overnight at 4 °C. Secondary antibody against mouse IgM (Cat. #ab175662, Abcam) was used to detect α-Ceramide antibody. Tom20 and Ceramide were imaged and quantified by a Nikon A1-Confocal.

Confocal images were acquired with a Nikon A1-Confocal using a Plan-Apo 60X 1.4 N.A. oil objective. Z-stacks were collected with a z-interval of 420 nm and pixel size of 100 nm using 2-frame Kallman-averaging. DAPI was excited with a 405 nm laser line; emission was collected via 450/50 bandpass filter. GPF was excited with a 488nm laser line; emission was collected via 525/42 bandpass filter. mCherry was excited at 561 nm and emission was collected via 595/50 filter. Detector gains, amplifier offsets and laser power were adjusted to maximize the range of signal for each channel. All images were collected and processed with NIS-Elements; deconvolution was performed using 20 interations. Widefield fluorescence images were acquired with a Zeiss Observer Z1 and Slidebook 6.0 software (3i) using a PlanApo 20X and a PlanApo 40X oil 1.3 NA.

#### Immunoblots

Antibodies include anti-PINK1 (Cell Signaling, 6946T), anti-Parkin-1 (Cell Signaling, 4211T), anti-NDP52 (Cell Signaling, 60732T), anti-SQSTM1 (Cell Signaling, 8025T), anti-BNIP3L/Nix (Cell Signaling, 12396T), anti-LC3B (Cell Signaling, 368T), anti-DRP1 (Abcam, ab56788), anti-DMN2 (Santa Cruz, Sc-166669), anti-MFN1 (Cell Signaling, 14739S), anti-MFN2 (Cell Signaling, 9482S), anti-OPA1 (Cell Signaling, 67589S), anti-GAPDH (Santa Cruz, Sc-32233), and anti-Beta-Actin (Santa Cruz, Sc-47778).

#### Isolation of Extracellular Vesicles by density gradient flotation

Extracellular vesicles were isolated as previously described.^20^ Briefly, large EVs were depleted from samples by centrifugation at 2000×*g* for 20 min and 16,500×*g* for 30min. Resulting supernatant was filtered through a 0.22 µm vacuum filter and was densified by mixing with OptiPrep iodixanol solution (Sigma D1556) to a final density of 1.16-1.30 g/mL, bottom-loaded into a polycarbonate ultracentrifuge tube and overlaid with 0.5-2mL aliquots of iodixanol/PBS solution in the 1.20-1.01 g/mL range to form a density gradient. Ultracentrifugation was performed for at 100,000×*g* for 4 hrs. Vesicles were collected from the top of the tube. Density of harvested fractions was measured based on iodixanol absorbance at 250 nm using a NanoDrop spectrophotometer (Thermo Fisher Scientific, Wilmington, DE). Vesicle harvests were stored at − 80°C.

#### Subcellular Fractionation and Isolation of Organelles

Cell culture media was removed by aspiration and the cells were washed twice with ice cold PBS, followed by washing and 5 min incubation in cold hypotonic lysis buffer (25 mM Tris·HCl, pH 7.5, 50 mM sucrose, 0.5 mM MgCl_2_, 0.2 mM EGTA). Buffer was removed by aspiration and the cells scraped with harvest volume adjusted to 4 mL with hypotonic buffer. Cells were lysed ten cycles through a ball-bearing homogenizer (isobiotec, Heidelberg, Germany); lysis was confirmed by microscopic examination. The cellular homogenate was made isotonic by addition of 356 µL of hypertonic buffer (25 mM Tris base, pH 7.2, 2.5 M sucrose, 0.5 mM MgCl_2_, 0.2 mM EGTA). Initial subcellular fractionation was performed by double serial centrifugation at 4°C at 300, 5,000 and 10,000x*g* to yield crude nuclear, mitochondrial, microsomal and cytosolic fractions. The crude compartments were further purified by centrifugation at 4°C at 10,000-50,000x*g* in discontinuous gradients of 10, 17.5, 25, 30 and 35% iodixanol in isotonic Tris-sucrose buffer (25 mM Tris base, pH 7.2, 2.5 M sucrose, 0.5 mM MgCl_2_, 0.2 mM EGTA) to yield nuclear, mitochondrial, plasma membrane, ER and heavy microsomal fractions as described by Andreyev, et al.^21^

### Untargeted Lipidomic Analyses

#### Sample Extraction

Cell lysates were washed 2x with pre-chilled 0.9% NaCl followed by addition of 2.5mL of pre-chilled 3:1 isopropanol:ultrapure water. Cells were scraped using a 25cm Cell Scraper (Sarstedt) in extraction solvent and transferred to a 15mL conical tube (Eppendorf). Samples were briefly vortexed followed by centrifugation at 4°C for 10 min at 2,000 x g. Thereafter, 1.2mL of metabolite extracts were transferred to 1.5mL Eppendorf tubes and stored in −20°C until lipidomic analyses. For analysis of complex lipids, in a 96 well plate, 10µL (3:1 isopropanol:ultrapure water) of the cell lysates supernatant was diluted with 90µL of 1:3:2 100mM ammonium formate, pH3: acetonitrile: 2-propanol (Fisher Scientific) and transferred to a 384-well microplate (Eppendorf) for analysis by LC-MS.

#### Untargeted Analysis of Complex Lipids

Untargeted lipidomic analyses were conducted on a Waters Acquity™ UPLC system coupled to a Xevo G2-XS quadrupole time-of-flight (qTOF) mass spectrometer as previously described.^22,23^ Chromatographic separation was performed using a C18 (Acquity™ UPLC HSS T3, 100 Å, 1.8 µm, 2.1×100mm, Water Corporation, Milford, U.S.A) column at 55°C. The mobile phases were (A) water, (B) Acetonitrile, (C) 2-propanol and (D) 500mM ammonium formate, pH 3. A starting elution gradient of 20% A, 30% B, 49% C and 1% D was increased linearly to 4% A,14% B, 81% C and 1 % D for 4.5 min, followed by isocratic elution at 4% A,14% B, 81%C and 1%D for 2.1 min and column equilibration with initial conditions for 1.4 min.

#### Mass Spectrometry Data Acquisition

Mass spectrometry data was acquired using ‘sensitivity’ mode in positive and negative electrospray ionization mode within 100-2000 Da. For the electrospray acquisition, the capillary voltage was set at 1.5 kV (positive), 3.0kV (negative), sample cone voltage 30V, source temperature at 120° C, cone gas flow 50 L/h and desolvation gas flow rate of 800 L/h with scan time of 0.5 sec in continuum mode. Leucine Enkephalin; 556.2771 Da (positive) and 554.2615 Da (negative) was used for lockspray correction and scans were performed at 0.5sec. The injection volume for each sample was 3 µL. The acquisition was carried out with instrument auto gain control to optimize instrument sensitivity over the samples acquisition time.

LC-MS and LC-MSe data were processed using Progenesis QI (Nonlinear, Waters. Annotations were determined by matching accurate mass and retention times using customized libraries created from authentic standards and/or by matching experimental tandem mass spectrometry data against the NIST MSMS or HMDB v3 theoretical fragmentations; for complex lipids retention time patterns characteristic for lipid subclasses are also considered.

To correct for injection order drift, each feature was normalized using data from repeat injections of quality control samples collected every 10 injections throughout the run sequence. Measurement data were smoothed by Locally Weighted Scatterplot Smoothing (LOESS) signal correction (QC-RLSC) as previously described.^22,23^ Feature values between quality control samples were interpolated by a cubic spline. Only detected features exhibiting a relative standard deviation (RSD) less than 30 in either the historical or pooled quality controls samples were considered for further statistical analysis. Metabolite values were normalized to the total ion count and reported as normalized area units.

### Proteomic Analyses

#### Proteomic Profiling of Breast Cancer Cell Lines

Twenty-eight Breast cancer cell lines MCF7, MDAMB231, SKBR3, HCC1954, HCC1143, BT474, HCC1500, T47D, ZR751, HCC1937, HCC1599, HCC202, HCC1806, MDAMB468, HCC2218, HCC70, HCC1187, Hs578T, BT549, MCF10A, MCF12A, MDAMB361, HCC1395, CAMA1, HCC38, MDAMB436, BT20 and MDAMB157 were labeled with 13C6 Lys (#CNLM-2247, Cambridge isotope laboratories) in RPMI1640 containing 10% dialyzed FBS and 1 % penicillin/streptomycin cocktail (Gibco). SILAC labeling was to discriminate FBS derived proteins from cell proteins.

#### Proteomic Profiling of Breast Cancer Cell Lines

For proteomic analysis of whole-cell lysates, −2 × 107 cells were lysed in 1 mL of PBS containing octyl-glucoside (1% w/v) and protease inhibitors (complete protease inhibitor cocktail, RocheDiagnostics), followed by sonication and centrifugation at 20,000 × g with collection of the supernatant, and filtration through a 0.22µm filter. Two milligrams of WCE proteins were reduced in DTT and alkylated with acrylamide before fractionation with RP-HPLC. A total of 84 fractions were collected at a rate of 3 fractions/min. Mobile phase A consisted of H2O:ACN (95:5, v/v) with 0.1% of TFA. Mobile phase B consisted of ACN:H2O (95:5) with 0.1% of TFA. Collected fractions from were dried by lyophilization followed by in-solution digestion with trypsin (Mass Spectrometry Grade, Thermo Fisher).

Based on the chromatogram profile, 84 fractions were pooled into 24 fractions for LC-MS/MS analysis per cell line. In total, 2,688 fractions were analyzed by RPLC-MS/MS using a nanoflow LC system (EASYnano HPLC system, Thermo Scientific) coupled online with LTQ Orbitrap ELITE mass spectrometer (Thermo Scientific). Separations were performed using 75 µm id × 360 µm od × 25-cm-long fused-silica capillary column (Column Technology) slurry packed with 3 µm, 100 A° pore size C18 silica-bonded stationary phase. Following injection of ∼2µg of protein digest onto a C18 trap column (Waters, 180 µm id×20 mm), peptides were eluted using a linear gradient of 0.35% mobile phase B (0.1 formic acid in ACN) per minute for 90min, then to 95% B in an additional 10 min, all at a constant flow rate of 300 nL/min. Eluted peptides were analyzed by LTQ Orbitrap ELITE in data-dependent acquisition mode. Each full MS scan (m/z 400–1800) was followed by 20 MS/MS scans (CID normalized collision energy of 35%). Acquisition of each full mass spectrum was followed by the acquisition of MS/MS spectra for the 20 most intense +2, +3 or +4 ions within a duty cycle, dynamic exclusion was enabled to minimize redundant selection of peptides previously selected for MS/MS analysis. Parameters for MS1 were 60,000 for resolution, 1 × 106 for automatic gain control target, and 150 ms for maximum injection time. MS/MS was done by CID fragmentation with 3×104 for automatic gain control, 10 ms for maximum injection time, 35 for normalized collision energy, 2.0 m/z for isolation width, 0.25 for activation q-value, and 10 ms for activation time.

MS/MS spectra were searched against the Uniprot proteome database (Human and Bovine, January 2017) using X!Tandem search engine through Trans-Proteomic Pipeline (TPP 4.8) and processed with the peptide and protein prophet. Trypsin was specified as protein cleavage site, with possibility of two missed cleavages allowed. For the modifications, one fixed modification of propionamide (71.037114) at Cysteine and two variable modifications, oxidation at methionine (15.9949 Da) and SILAC 13C6 at lysine (6.0201 Da) were chosen. Addition of SILAC was to strictly discriminate human protein with bovine proteins and not intended to perform relative quantitation of Heavy versus Light ratios. The mass error allowed was 10 ppm for parent monoisotopic and 0.5 Da for MS2 fragment monoisotopic ions. The searched result was filtered with FDR=0.01.

#### Proteomic Profiling of Extracellular Vesicles

EV-derived protein digestion and identification by LC-MS/MS was performed using established protocols.^22^ NanoAcquity UPLC system coupled in-line with WATERS SYNAPT G2-Si mass spectrometer was used for the separation of pooled digested protein fractions. The system was equipped with a Waters Symmetry C18 nanoAcquity trap-column (180 μm × 20 mm, 5 μm) and a Waters HSS-T3 C18 nanoAcquity analytical column (75 μm × 150 mm, 1.8 μm). The column oven temperature was set at 50 °C, and the temperature of the tray compartment in the auto-sampler was set at 6 °C. LC-HDMSE data were acquired in resolution mode with SYNAPT G2-Si using Waters Masslynx (version 4.1, SCN 851). The capillary voltage was set to 2.80 kV, sampling cone voltage to 30 V, source offset to 30 V and source temperature to 100 °C. Mobility utilized high-purity N2 as the drift gas in the IMS TriWave cell. Pressures in the helium cell, Trap cell, IMS TriWave cell and Transfer cell were 4.50 mbar, 2.47 × 10−2, 2.90, and 2.53 × 10−3 mbar, respectively. IMS wave velocity was 600 m s−1, helium cell DC 50 V, Trap DC bias 45 V, IMS TriWave DC bias V and IMS wave delay 1000 μs. The mass spectrometer was operated in V-mode with a typical resolving power of at least 20,000. All analyses were performed using positive mode ESI using a NanoLockSpray source. The lock mass channel was sampled every 60 s. The mass spectrometer was calibrated with a [Glu1] fibrinopeptide solution (300 fmol µL−1) delivered through the reference sprayer of the NanoLockSpray source.

LC-HDMSE data were acquired in resolution mode with SYNAPT G2-Si using Waters Masslynx (version 4.1, SCN 851). The acquired LC-HDMSE data were processed and searched against protein knowledge database (Uniprot) through ProteinLynx Global Server (PLGS, Waters Company) with 4% FDR. The modification search settings included cysteine (Cys) alkylation with acrylamide (71.03714@C) as a fixed modification, and methionine (Met) oxidation (15.99491@M), acetylaldehyde (AA) (43.01839@K), malondialdehyde-acetylaldehyde (MAA) (136.05243@K), malondialdehyde (MDA) (55.01839@K) as a variable modification. The searched data was filtered with False Discovery Rate 4%.

### Gene expression profiles

Gene expression data for breast cancer cell lines were obtained from Cancer Cell Line Encyclopedia (CCLE) (www.broadinstitute.org/ccle).

### Human Cohort

Human plasma samples were obtained with informed consent and the studies were conducted under protocols approved and supervised by the University of Texas MD Anderson IRB (LAB03-0479). The cohort consisted of a case-control cohort comprised of 85 TNBC breast cancer cases and 141 cancer-free (minimum of 3-year follow-up) controls.^24^

### BRCA1^co/co^; MMTV-Cre; p53^+/-^ orthotopic syngeneic mouse TNBC cancer model

All animal experiments were conducted in accordance with accepted standards of humane animal care approved by MD Anderson Cancer Center (MDACC) IACUC. BRCA1^co/co^; MMTV-Cre; p53^+/-^ mouse TNBC cancer cells, transduced with firefly lentiviral particle (Firefly Luciferase Lentifect^TM^, GeneCopoeia, MD) and selected by puromycin with 5 μg/ml dose concentration, were injected into 8-10 weeks old B6129SF1/J mice (Jackson Labs, Maine, Stock No.101043).

For initial testing, luciferase labeled BRCA1^co/co^; MMTV-Cre; p53^+/-^-Luc TNBC cancer cells (1 × 10^6^ in 50μL (1:10, Matrigel:HBSS)) were injected into the mammary fat pad of B6129SF1/J mice. Ten days after cell injection, tumor growth was assessed using Xenogen IVIS-200 bioluminescence imaging (PerkinElmer) and mice were randomized into treatment groups: (*n*=7) daily oral gavage of 60 mg kg^−1^ of eliglustat (AbMole BioScience, M9733) or (*n*=7) saline control. Both groups of tumor-bearing mice received eliglustat or saline for 14 days, after which tumors and blood samples were collected for further analysis. Mice were monitored for signs of physical distress and changes in body weight throughout the intervention. For dose de-escalation studies, BRCA1^co/co^; MMTV-Cre; p53^+/-^ tumor bearing mice were treated via daily oral gavage with 15, 30, and 60 mg kg^−1^ of eliglustat (*n*=4 mice per group) or saline control (*n*=4 mice) for 14 days. For survival analyses, an independent cohort of BRCA1^co/co^; MMTV-Cre; p53^+/-^ tumor bearing mice were treated via daily oral gavage with 30 mg kg^−1^ of eliglustat (*n*=3 mice per group) or saline control (*n*=3 mice). Tumor size and body weight were measured and recorded twice a week. Tumor volume was calculated by the formulations V = (L x W x W)/2, where V is tumor volume, L is tumor length, W is tumor width.

### Immunohistochemistry

Primary antibodies to Ki67 (catalog #ab16667, Abcam, 1:500 dilution), Caspase-3 (catalog #ab4051, Abcam, 1:200 dilution), LC3B (catalog #ab48394, Abcam, 1:2000 dilution), and PINK1 (catalog #ab23707, Abcam, 1:1000) were used for immunohistochemistry analysis on formalin-fixed paraffin-embedded tissue slides from the BRCA1^co/co^; MMTV-Cre; p53^+/-^ mouse TNBC tumors. Sections were deparaffinized in xylene, rehydrated in a descending ethanol series, and then treated with 3% hydrogen peroxide for 20 minutes. Antigen retrieval was conducted in a pressure cooker in 1X ImmunoRetriever with Citrate pH 6.0 (Bio SB, Santa Barbara, California, USA) and 0.1% Tween 20 at 121°C for 15 minutes. After a 30-minute incubation in 0.5% casein milk in TBST, tissue sections were incubated in primary antibodies for 16 hours at 4°C, followed by incubation in HRP rabbit polymer conjugated secondary antibody (Cell Signaling, catalog#8114S). The immunoreaction was visualized in Histofine DAB-2V Kit (Nichirei Bioscience, Tokyo, Japan, catalog#425314F) following the manufacturer’s procedure. Images were scanned by MDACC-Histology Core Laboratory and analyzed using Aperio Imagescope (Leica Biosystems, Buffalo Grove, Illinois, USA).

### Blood chemistry tests and Tissue Anatomic Pathology

Blood chemistry tests and histopathological examination of liver and kidney tissues from eliglustat treated BRCA1^co/co^; MMTV-Cre; p53^+/-^ TNBC-tumor bearing mice were performed under established protocols by the UTMDACC-DVMS Clinical Pathology Lab.

### Statistical Analyses

Statistical significance was determined using Student t-tests or Wilcoxon rank sum tests. For comparison of metabolic profiles of isolated mitochondria following eliglustat or vehicle control, paired t-test was used. Statistical significance was determined at p<0.05 and all p-values are reported as 2-sided unless otherwise specified. Figures were generated in Prism Version 8 (GraphPad Software, Inc).

## Results

### Mitochondrial dynamics in TNBC cells

Evaluation of whole cell lysate proteomic profiles from a panel of 28 breast cancer cell lines, comprising TNBC (n=17) and non-TNBC (n=11) subtypes, revealed expression of mitochondrial fusion- and fission-related proteins to be similar between TNBC and non-TNBC cell lines, with exception of the pro-fission protein dynamin 2 (DNM2), which was statistically significantly higher in TNBC cells (Wilcoxon Rank Sum Test 2-sided P<0.05) (**Fig. 1A; Table S1**). Mitophagy-related proteins Optineurin (OPTN) and LC3B-II were also found to exhibit significantly (Wilcoxon Rank Sum Test P<0.05) higher expression in TNBC compared to non-TNBC cell lines (**Fig. 1A; Table S1**). Western blot analyses and CCLE-derived gene expression analyses provided concordant results (**Fig. S1-S2**). Moreover, as exemplified in the MDAMB231 TNBC cell line, TNBC cells evidenced heterogenous mitochondrial architectures with some cells displaying a highly-networked (*fused*) mitochondrial architecture (see cells 1 and 2 of **Fig. 1B** labeled with mitochondrial marker PDHA-RFP CellLight construct), in contrast to other cells exhibiting a more punctate mitochondrial architecture, with discrete mitochondria observed to be co-localized with the autophagosome/lysosome, suggesting an active state of mitophagy (see cells 3 and 4 of **Fig. 1B** labeled with PDHA-RFP and autophagosome/lysosomal marker LAMP1-GFP CellLight constructs).

**Figure 1.**
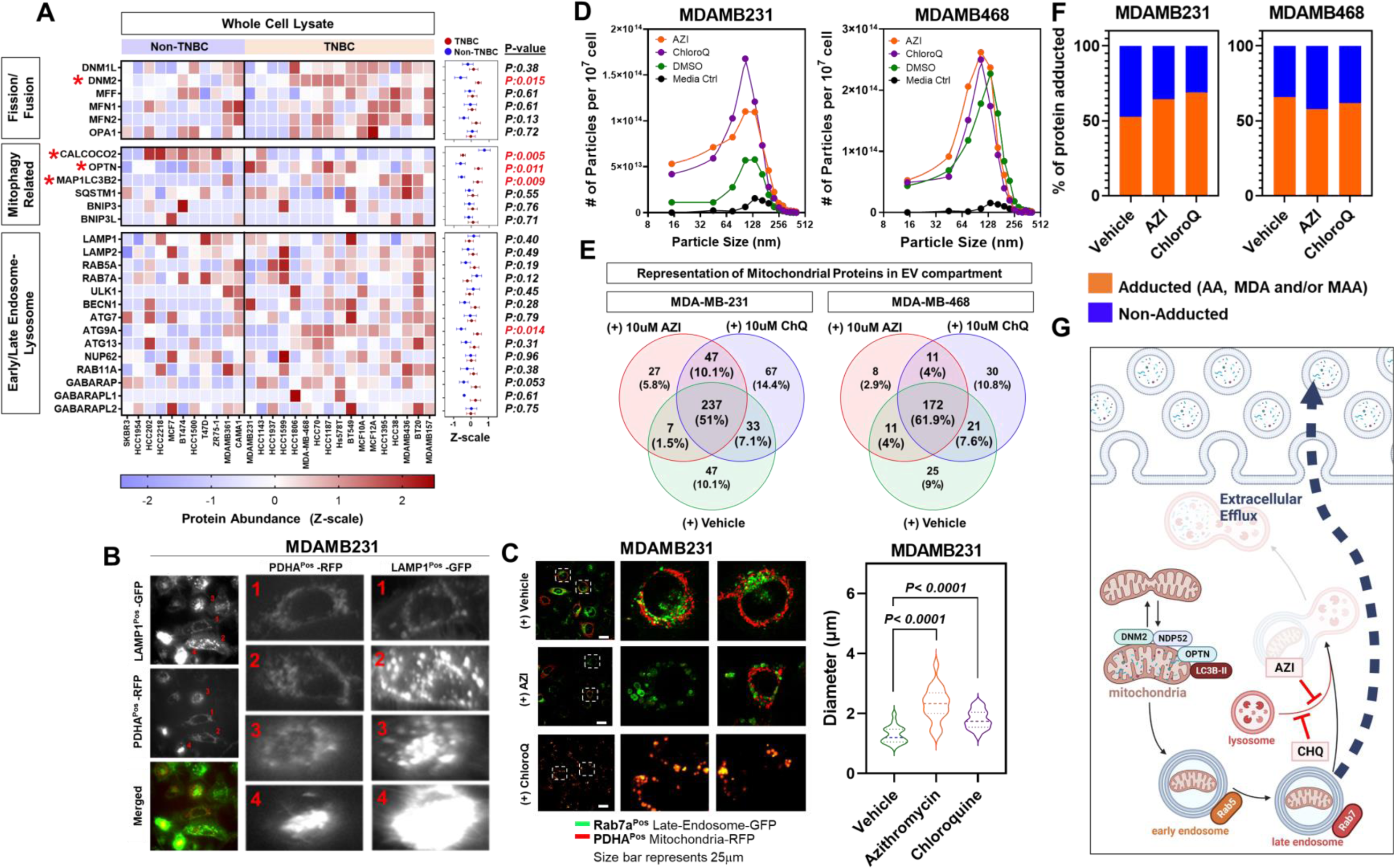
Intersection between cancer-disseminated EVs and mitochondrial dynamics. **A)** Heatmap depicting protein expression (z-scores) for proteins involved in mitochondrial fission/fusion, mitophagy, and early- and late-endosome and endolysosome across 28 breast cancer cell lines. Dot plot to the right of the heatmaps shows mean +/- standard deviation of z-scores for TNBC (red nodes) and non-TNBC (blue nodes) cell lines. Statistical significance was determined by 2-sided Wilcoxon Rank Sum Test. **B)** Staining for mitochondria (CellLight Mitochondria-RFP (leader sequence of E1 alpha pyruvate dehydrogenase) and lysosomes (CellLight Lysosome-GFP (lysosomal associated membrane protein 1)) in MDAMB231 TNBC cells. Individual panels represent subpopulations of cancer cells exhibiting a more ‘fused’-like mitochondrial architecture whereas other cell subpopulations exhibited punctate, ‘hyperfused’ mitochondria that were co-localized with the lysosome. **C)** Staining for mitochondria (CellLight Mitochondria-RFP) and late-endosome (CellLight Ras-related protein Rab-7a (Rab7a)-GFP) in MDAMB231 TNBC cells treated with either vehicle (DMSO) control or 10uM Azithromycin (AZI). Enlarged images highlight ‘swelling’ of late endosome following treatment of AZI. Quantitation of mean diameter of individual endosomes are provided in violin plots to the right. **D)** Particle counts per mL of conditioned media following 48-hour treatment with either vehicle (DMSO) control, 10uM AZI, or 10uM chloroquine. Control media is included as a reference control. **E)** Venn diagram illustrating overlap between proteins quantified in extracellular vesicles (EVs) and isolated mitochondria from MDAMB468 and MDAMB231 TNBC cell lines. **F)** Bar plots illustrating proportion of proteins identified in EVs that have an acetaldehyde (AA), malondialdehyde (MDA), and malondialdehyde-acetaldehyde (MAA) adduct. **G)** Schematic depicting intersection of EV biogenesis and mitochondrial clearance.

To determine if mitophagic flux was active in TNBC cells, we treated TNBC cells with the autophagy/mitophagy inhibitors azithromycin (AZI), bafilomycin, or chloroquine (CHQ) for 24 hours and performed immunoblot analyses for mitophagy related protein markers. Inhibitor-treated TNBC cells showed pronounced increases in protein expression of SQSTM1/p62 and LC3BII (**Fig. S3**). Functionally, p62/SQSTM1 acts as a selective autophagy receptor that targets cargoes for specific degradation in the proteosome. LC3BII recruits p62 attached to select ubiquitinated cargo molecules for engulfment into the autophagosome and their subsequent delivery to the lysosome for degradation. During this process, both p62 and LC3BII are degraded.^25–30^ Thus, the observed accumulation of p62 and LC3BII following treatment with autophagy/mitophagy inhibitors is consistent with baseline active mitophagy in TNBC cells. In support of this, treatment of MDAMB231 TNBC cells with AZI or CHQ resulted in enlargement of late endosomes (as indicated by labeling with late endosomal marker Rab7A-GFP CellLight construct; violin plots depicting mean diameter of individual endosomes are provided to the right) coincident with mitochondrial components (as indicated by labeling with PDHA-RFP CellLight construct (**Fig. 1C**)), suggesting failure of autophagosome fusion with the lysosome.^31^

Notably, the above-mentioned changes were met with concurrent increases in cancer cell disseminated EVs into cell conditioned medium (CM) (**Fig. 1D**). We considered that the increased cancer cell secretion of EVs may manifest an alternative route for removing mitochondrial damage. To test this hypothesis, we first determined whether cancer cell-derived EVs contained mitochondrial proteins by comparing proteomic profiles generated on isolate mitochondria (**Fig. S6**) and CM-derived EVs from MDAMB231 and MDAMB468 TNBC cells cultured in lipid-free FBS containing media under basal conditions and following 24-hour treatment with AZI or CHQ. Proteomic analyses revealed a high representation of mitochondrial proteins in cancer-disseminated EVs (**Fig. 1E; Table S2-3**). Moreover, several endosomal and endo-lysosomal markers, including RAS-Associated Protein RAB5A (Rab5A) and Rab7A, lysosomal associated membrane protein 1 (LAMP1) and LAMP2, as well as mitophagy-related proteins including p62/SQSTM1 and LC3B were detected in cancer cell disseminated EVs (**Table S3**).^32–34^

Elevated mitochondrial stress promotes formation of reactive aldehydes including acetaldehyde (AA), malondialdehyde (MDA), and malondialdehyde-acetaldehyde (MAA) that may lead to protein-adducts and impaired protein function.^35,36^ Leveraging EV proteomic profiles, we additionally evaluated for aldehyde protein adduction. These analyses revealed >50% of EV-associated proteins to be aldehyde adducted (**Fig. 1F**). Collectively, the above findings suggest an alternative route by which cancer cells can remove ‘damaged’ mitochondrial components via EVs (**Fig. 1G**). These findings are consistent with a recent study demonstrating that mouse embryonic fibroblasts remove mitochondria via EVs when lysosomal function is compromised.^11^

### TNBC cells scavenge extracellular sphingomyelins to sustain elevated glycosphingolipid biosynthesis to support EV biogenesis

We next aimed to define the onco-metabolic processes and pathways that support EV-mediated clearance of mitochondria. We investigated cell lipid metabolism based on the composition of EVs as lipid bilayer entities that traffic protein and other materials ^37^, as well as our prior findings of lipidomic alterations in prostate cancer.^22^

We initially assessed whether restricting extracellular lipid availability would impact sensitivity of TNBC cells to autophagic/mitophagic inhibitors. Compared to normal culture conditions (10% FBS), TNBC cells exhibited increased sensitivity to CHQ and AZI when grown in low serum (1%) or lipid-free serum (**Fig. 2A**). Inversely, the anti-cancer effects of CHQ and AZI were partially mitigated by media supplementation with synthetic self-assembled lipid particles (SSALPs) (**Fig. 2B**; **Fig. S3**). These findings suggest a process of cellular scavenging of extracellular lipids to support EV-mediated clearance of damaged mitochondria.

**Figure 2.**
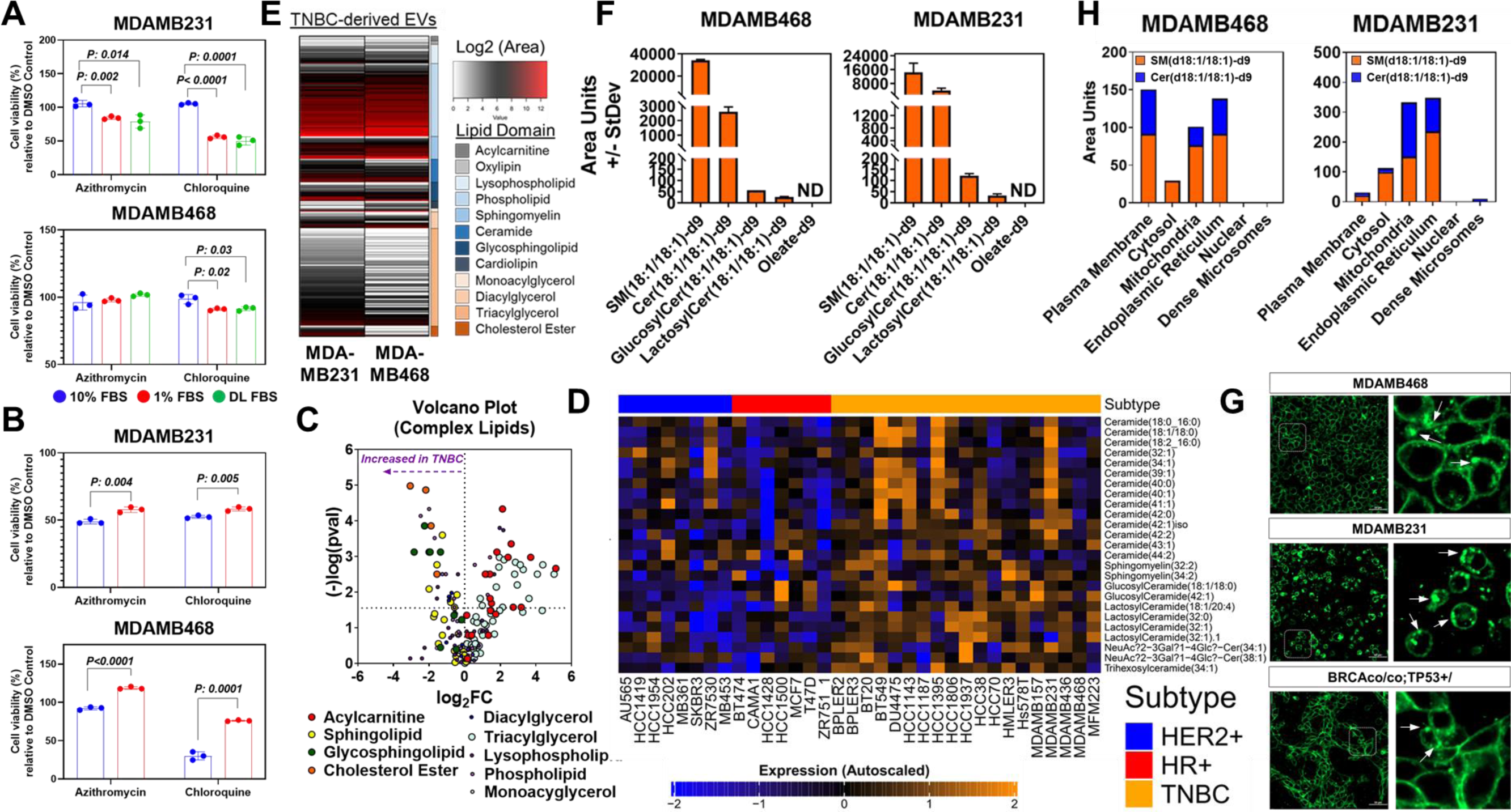
TNBC salvage extracellular sphingomyelin to sustain elevated glycosphingolipid biosynthesis. **A)** % viability (MTS assay) relative to vehicle (DMSO) control for MDAMB231 and MDAMB468 TNBC cells following 48 hour treatment with 10uM azithromycin (AZI) or chloroquine (CHQ) in the presence of 10% FBS, 1% FBS, or 10% delipidated FBS. Statistical significance was determined by 2-sided Student T-test. **B)** % viability (MTS assay) relative to vehicle (DMSO) control for MDAMB231 and MDAMB468 TNBC cells following 48 hour treatment with 10uM AZI or CHQ in medium supplemental with or without self-assembled lipid particles (SSALPs). Composition of SSALPs is provided in **Supplemental Figure S3.** Statistical significance was determined by 2-sided Student T-test. **C)** Volcano plot illustrating differences in lipid composition between TNBC and non-TNBC cell lines. Nodes represent individual lipid species belonging to the respective lipid sub-class. **D)** Heatmap illustrating differences in whole cell lysate levels of quantified sphingolipids and glycosphingolipids among 34 breast cancer cell lines. **E)** Lipidomic profiles of EVs isolated from conditioned medium of MDAMB231 and MDAMB468 TNBC cell lines. **F)** Relative abundance (area units +/- StDev) of sphingomyelin(18:1/18:1)-deuterium(d)9 as well as ceramide(18:1/18:1)-d9, glucosylceramide((18:1/18:1)-d9, lactosylceramide((18:1/18:1)-d9, and oleate-d9 isotopologues in whole cell lysates of TNBC cell lines MDAMB468 and MDAMB231 following 48 hour treatment with self-assembleded lipid particles (SSALPs) enriched in sphingomyelin(18:1/18:1)-d9. **G)** Uptake of C11 TopFluor-SM in MDAMB468 and MDAMB231 TNBC cell lines as well as murine BRCAco/co/TP53+/- TNBC cell line. White arrows illustrate subcellular aggregation of C11 TopFluor-SM. **H)** Relative abundances of sphingomyelin(18:1/18:1)-d9 and the ceramide(18:1/18:1)-d9 isotopologue in different subcellular compartments of MDAMB468 and MDAMB231 TNBC cell lines following 48 hour treatment with SSALPs enriched in sphingomyelin(18:1/18:1)-d9.

Next, leveraging lipidomic data for a panel of 34 breast cancer cell lines (7 hormone receptor positive (HR+), 8 HER2-enriched, and 19 TNBC cell lines) (**Table S4**). Compared to non-TNBC, 37 lipid species that were statistically significantly (Wilcoxon rank sum test 2-sided p<0.05) elevated in TNBC cell lines whereas 36 lipid species were reduced (**Table S4**). Among TNBC-enriched (Wilcoxon rank sum test 2-sided p<0.05) lipids were several sphingolipids (**Fig. 2C-D**). Of relevance, lipidomic analyses of EVs isolated from CM of TNBC cells also revealed enrichment of sphingolipids and glycated derivatives (**Fig. 2E; Table S4**). We therefore considered that the heighted intracellular levels of sphingolipids in TNBC cells may thus support biogenesis of (glyco)sphingolipid-enriched EVs.

Ceramides serve as the backbone for glycosphingolipids and are derived through three primary metabolic pathways: *de novo*, recycling of sphingosine, and a salvaging pathway via hydrolysis of sphingomyelins (SMs) by sphingomyelinases (**Fig. S4A**). To determine the source of ceramides, we first evaluated protein expression of ceramide-associated enzymes using the proteomic profiles from the 28 TNBC cell lines. These analyses revealed protein levels of delta 4-desaturase, sphingolipid 2 (DEGS2), and ceramide synthase 2 (CERS2), enzymes associated with de novo and sphingosine recycling pathways, respectively, to be statistically significantly (Wilcoxon rank sum test 2-sided p<0.05) lower in TNBC compared to non-TNBC cell lines (**Fig. S4B-C; Table S5**). Sphingomyelinases, particularly SPMD4, tended to be elevated in TNBC compared to non-TNBC cell lines (**Fig. S4B**), implicating sphingomyelin salvaging as a potential source of ceramides and glycosphingolipid derivatives. To test this hypothesis, we treated TNBC cells for 48 hours with SSALPs containing deuterated SM(18:1/18:1)-d9 and traced its biochemical fate using liquid chromatography mass spectrometry (LCMS). In addition to SM(18:1/18:1)-d9, we detected the ceramide(18:1/18:1)-d9, glucosylceramide(18:1/18:1)-d9, and lactosylceramide(18:1/18:1)-d9 isotopologues (**Fig. 2F**), thereby confirming activity of the sphingomyelin salvaging pathway. Notably, oleate-d9 was not detected, indicating preferential shunting of ceramides into the glycosphingolipid pathway. Treatment and live cell confocal imaging of human MDAMB468 and MDAMB231 TNBC cells as well as murine BRCA1^co/co^; MMTV-Cre; p53^+/-^ TNBC cells with C11 TopFluor-SM further confirmed that TNBC cells can scavenge extracellular SM that was observed aggregated in discrete subcellular organelles (as indicated by white arrows in **Fig. 2G**). Subcellular organelle fractionation and lipidomic analyses (**Fig. S5**) following treatment of MDAMB468 and MDAMB231 TNBC cells with SM(d18:1/18:1)-d9 containing SSALPs further revealed mitochondria to be a predominant site for trafficking of SM(d18:1/18:1)-d9 as well as the ceramide(18:1/18:1)-d9 isotopologue (**Fig. 2H**).

Based on the above findings, we hypothesized that the scavenging of extracellular SM and subsequent metabolism to ceramide and its glycated derivatives may accompany biogenesis of (glyco)sphingolipid-enriched EVs (**Fig. 2E**) that contain jettisoned mitochondrial components (**Fig. 1E-F**).^22^ To test this, we treated TNBC cell lines with SM(18:1/18:1)-d9 containing SSALPs and performed subsequent LCMS analyses on CM, the results of which demonstrated accumulation of the ceramide(18:1/18:1)-d9, glucosylceramide(18:1/18:1)-d9 and hexosylceramide(18:1/18:1)-d9 isotopologues (**Fig. 3A**). Next, we perturbed mitochondrial homeostasis via short-term treatment with carbonyl cyanide m-chlorophenyl hydrazone (CCCP) and carbonyl cyanide-p-trifluoromethoxyphenylhydrazone (FCCP)^38^, and performed lipidomic analyses of isolated mitochondria and corresponding conditioned media. These studies revealed that CCCP- and FCCP-induced mitochondrial damage promotes accumulation of sphingomyelin and glycosphingolipids in isolated mitochondria that was met with paralleled increases in (glyco)sphingolipid-enriched EVs in conditioned media (**Fig. 3B; Fig. S6**). In contrast, deprivation of extracellular sphingomyelin bioavailability resulted in accumulation of punctate *hyperfused* mitochondria in MDAMB231 and MDAMB468 TNBC cells (**Fig. 3C**) and reduced cell invasiveness (**Fig. S5**).

**Figure 3.**
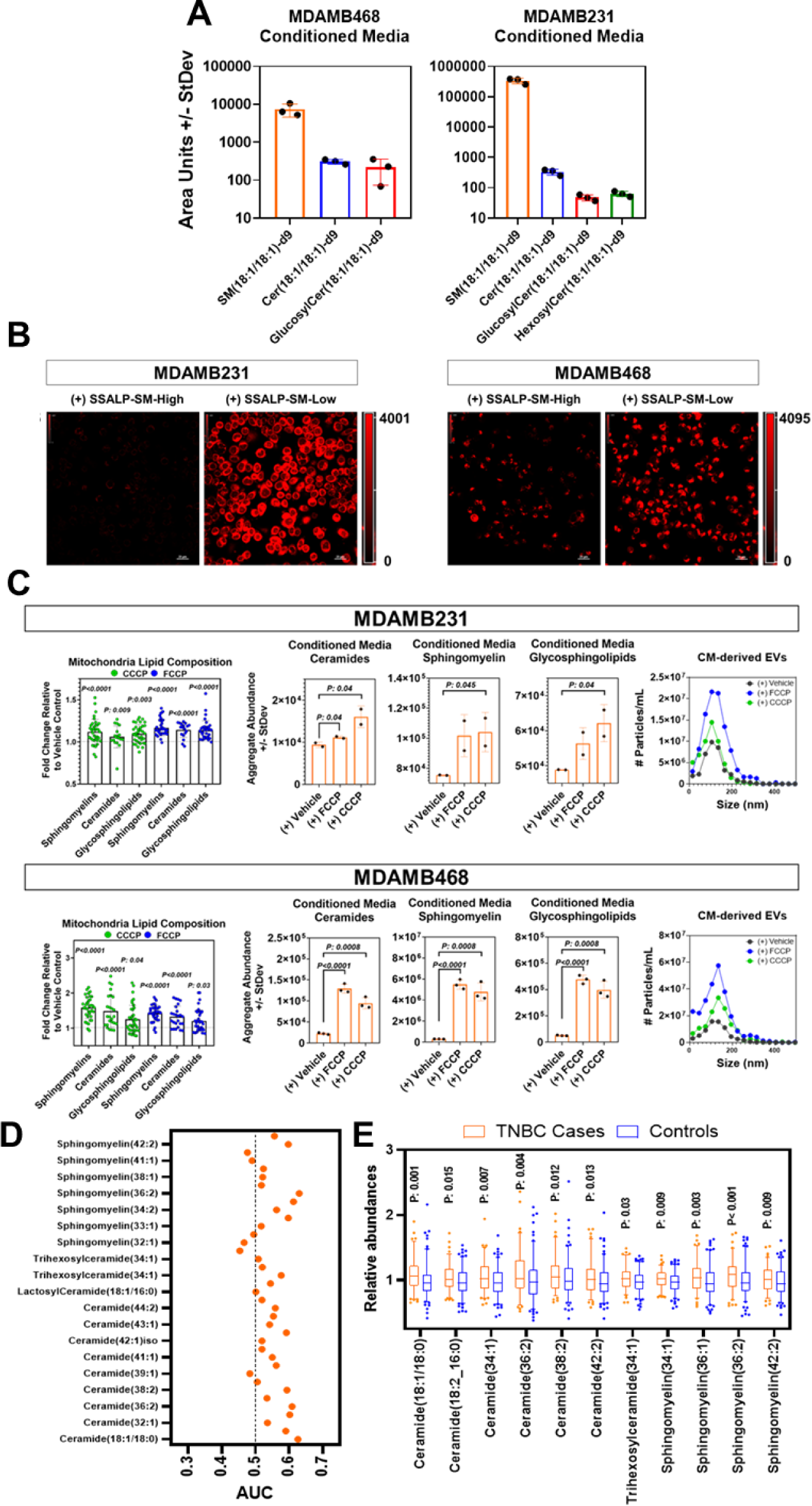
Extracellular vesicles enriched in (glycol)sphingolipids are essential for clearance of damaged mitochondria. **A)** Spectral abundance (area units) +/- StDev of SM(18:1/18:1)-d9 as well as ceramide(18:1/18:1)-d9, glucosylceramide(18:1/18:1)-d9, and hexosylceramide(18:1/18:1)-d9 in conditioned medium of MDAMB468 and MDAMB231 TNBC cells treated with SM(18:1/18:1)-d9 containing self-assembled lipid particles (SSALPs). **B)** Staining for mitochondria using MitoTracker Red CMXROS) in MDAMB231 and MDAMB468 TNBC cells following 48hr treatment with SSALPs containing high- or low-SM content (see methods). **C)** Abundances of sphingomyelin, ceramides, and glycosphingolipids in conditioned medium and isolated mitochondria of MDAMB231 and MDAMB468 TNBC cells following 6-hour treatment with carbonyl cyanide m-chlorophenyl hydrazone (CCCP) and carbonyl cyanide-p-trifluoromethoxyphenylhydrazone (FCCP). Particle counts per mL of conditioned media following treatment from an independent experiment is also provided. For mitochondria, each node represents a unique lipid belonging to the respective lipid class; values represent fold-change relative to vehicle control. For conditioned medium, values represent aggregate abundances of ceramides, sphingomyelin, or glycosphingolipids. P-values were determined by 1-sided Student T-tests. **D)** Dot plot illustrating area under the Receiver Operating Characteristics Curves of individual (glco)sphingolipids for distinguishing TNBC cases (n = 85) from cancer-free controls (n = 141). **E)** Box and whisker plots depicting distribution of (glyco)sphingolipids between TNBC cases and cancer-free controls. Statistical significance was determined by 2-sided Wilcoxon Rank Sum test.

Notably, lipidomic analyses of plasmas collected from 85 newly diagnosed TNBC cases and 141 cancer-free controls similarly revealed sphingolipid species to be statistically significantly (Wilcoxon Rank Sum Test 2-sided p<0.05) elevated in TNBC cases (**Fig. 3E-F**). Collectively, these findings support that elevated sphingomyelin uptake and subsequent salvaging to ceramide and glycosphingolipid derivatives is a metabolic adaptation of TNBC cells that regulates mitochondrial dynamics with pro-tumor effect.

### Inhibition of glucosylceramide synthase promotes mitochondrial lethality in BrCa cells

We tested the hypothesis that specific targeting of ceramide to glycosphingolipid conversion may present an actionable metabolic vulnerability in TNBC. We evaluated the efficacy of eliglustat, an selective inhibitor of glucosylceramide synthase (also known as UGCG; UDP-glucose:ceramide glucosyltransferase), a rate-limiting enzyme in glycosphingolipid metabolism, for effects on viability of TNBC cells *in vitro*. Treatment of 6 TNBC cell lines for 48 hours with eliglustat resulted in dose-dependent reductions in cell viability (**Fig. 4A**). Similar findings were observed in non-TNBC cell lines (**Fig. S7A**).

**Figure 4.**
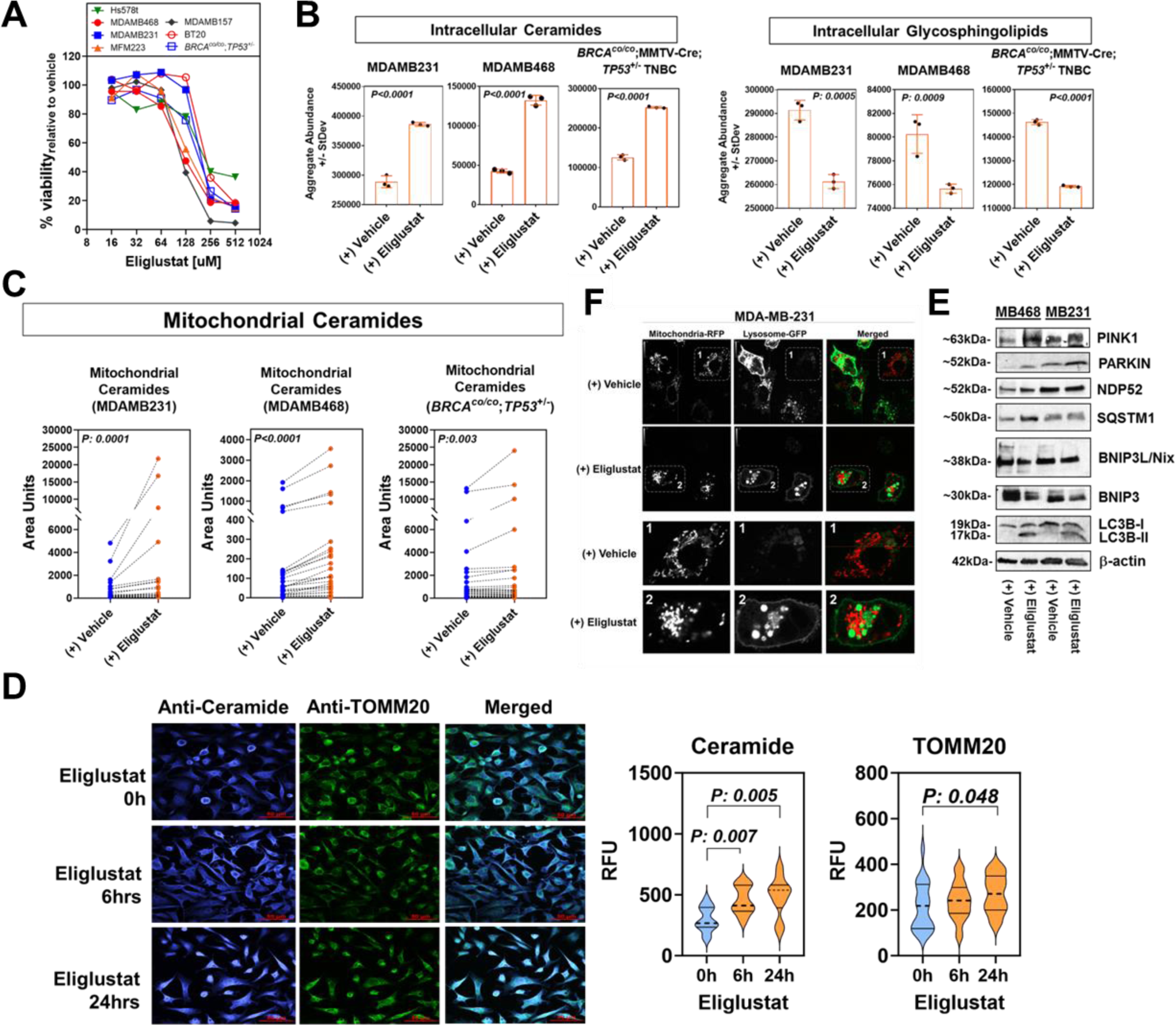
Small molecule inhibition of glucosylceramide synthase via eliglustat induces ceramide-associated mitochondria lethality in TNBC cells. **A)** Viability curves for a panel of 7 TNBC cell lines following treatment with eliglustat. Viability was determined by MTS assay. **B)** whole cell lysate levels of ceramides (left panel) and glycosphingolipids (right panel) in human MDAMB231 and MDAMB468 TNBC cells and murine BRCA1^co/co^; MMTV-Cre; p53^+/-^ TNBC cells following 6-hour eliglustat treatment. Values represent aggregate area abundances of ceramides or glycosphingolipids detected by mass spectrometry. Statistical significance was determined by 1-sided Student T-tests. **C)** Abundances (area units) of individual ceramide species detected by mass spectrometry in mitochondria isolated from human MDAMB231 and MDAMB468 TNBC cells and murine BRCA1^co/co^; MMTV-Cre; p53^+/-^ TNBC cells following 6-hour eliglustat treatment. Each node represents a unique ceramide; connecting lines show differences in area units for respective ceramide between vehicle and eliglustat treatment. Statistical significance was determined by 1-sided non-parametric paired t-test. **D)** Immunofluorescent staining of Tom20 (Green, Clone 1D6F5, Proteintech) and ceramide (Blue, Clone MID 15B4, Sigma-Aldrich) in MDAMB231 TNBC cells following treatment with 32μM eliglustat or vehicle control. All images were captured via confocal and widefield microscopy using live cell imaging techniques. Scale bars represent 50 µm; digital image acquisition parameters and look up table mappings were uniformly set for all confocal images. Quantitative values (relative mean intensities) are provided to the right of the images. **E)** Immunoblots for mitophagy-related proteins following 6-hour treatment with eliglustat or vehicle control in MDAMB231 and MDAMB468 TNBC cells. **F)** Staining for mitochondria (CellLight Mitochondria-RFP (leader sequence of E1 alpha pyruvate dehydrogenase) and lysosomes (CellLight Lysosome-GFP (lysosomal associated membrane protein 1)) in MDAMB231 TNBC cells following 6-hour treatment with eliglustat or vehicle control.

Lipidomics analyses of human MDAMB231 and MDAMB468 TNBC cells and murine BRCA1^co/co^; MMTV-Cre; p53^+/-^ TNBC cells^39^ following acute 6-hour treatment revealed statistically significant elevations in intracellular ceramide pools and reductions in glycosphingolipids (**Fig. 4B; Fig. S7B**). Subcellular analyses further revealed that eliglustat directly targeted the mitochondria that was accompanied by accumulation of ceramides ^40^ (**Fig. 4C-D; Fig. S8**), time-dependent increases in mitochondrial mass, as evidenced by increases in TOM20 staining (**Fig. 4D**), and elevated PINK1, PARKIN, and LC3B protein expression (**Fig. 4E**), suggesting induction of mitophagy. In support, assessment of mitochondrial morphology in MDAMB231 TNBC cells following acute (6hr) treatment with eliglustat indicated loss of the branched, fused-like-mitochondrial architecture with a shift towards accumulation of punctate ‘hyperfused’ mitochondria co-localized with the autophagosome/lysosome, suggesting induction of ‘lethal’ mitophagy (**Fig. 4F**).

### Eliglustat attenuated TNBC tumor growth in vivo

To evaluate the efficacy of eliglustat *in vivo*, we implanted BRCA1^co/co^; MMTV-Cre; p53^+/-^ TNBC-luciferase cells ^39^ into the mammary fat pad of B6129SF1/J mice. Tumor-bearing mice were initially treated daily via oral gavage with either saline control or 60mg/kg eliglustat (**Fig. 5A**).^22^ Tumor growth was suppressed by eliglustat (**Fig. 5B-C**). Immunohistochemical analyses showed that Ki67 staining was statistically significantly reduced in tumors following eliglustat treatment and caspase-3 staining positivity increased (**Fig. 5D**). Consistent with our *in vitro* studies, eliglustat-treatment increased protein expression of LC3B and PINK1 indicating induction of mitophagy (**Fig. 5D**). Notably, lipidomic analyses of plasma collected from BRCA1^co/co^; MMTV-Cre; p53^+/-^ TNBC tumor-bearing mice following eliglustat treatment revealed elevations in plasma cardiolipins, a mitochondria-exclusive lipid class known to be a key regulator of mitophagy (**Fig. 5E**).^41,42^ Of relevance, the elevation in plasma cardiolipins was concordant with the extent of tumor reduction following eliglustat challenge (**Fig. S9**). Thus, plasma cardiolipins appear to serve as a useful biomarker for monitoring the anti-cancer efficacy of eliglustat.

**Figure 5.**
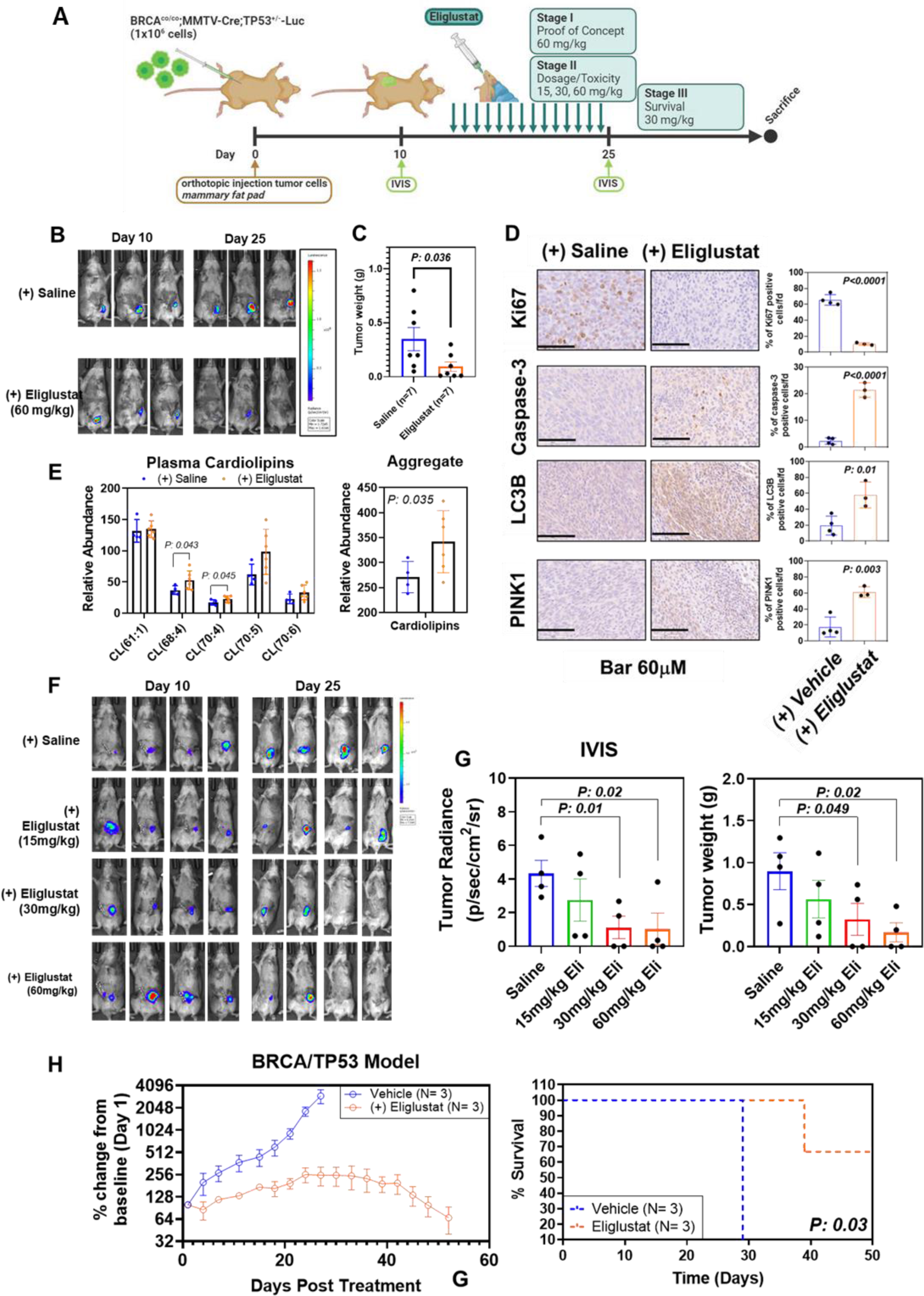
Eliglustat attenuated tumor growth and potentiates an anti-cancer immune response in a BRCA1^co/co^; MMTV-Cre; p53^+/-^ mouse model of TNBC. **A)** Schematic of treatment strategy for syngeneic BRCA1^co/co^; MMTV-Cre; p53^+/-^ mouse model of TNBC. Three phases were executed: 1- an initial pilot trial using 60mg/kg eliglustat; 2- a dose-escalation study (15, 30, and 60mg/kg); 3- prolonged treatment for survival analyses. **B)** Representative IVIS images following treatment. **C)** Tumor weight (g) following eliglustat treatment. Statistical significance was determined by 2-sided Wilcoxon rank sum test. **D)** Representative immunohistochemistry staining for Ki67, caspase-3, LC3B, and PINK1 in BRCA1^co/co^; MMTV-Cre; p53^+/-^ tumors following treatment with either saline or eliglustat (n=3-4 mice per group). Quantitative analyses (mean ± StDev) for respective markers are provided to the right of each representative IHC section. Scale bar represents 60 µM. Statistical significance was determined by 2-sided Student T-tests. **E)** Abundances of individual cardiolipins as well as aggregate abundances in plasma taken at the terminal time point from saline- or eliglustat-treated mice (n=4-7 mice per group). Statistical significance was determined 1-sided Student T-tests. **F)** Representative IVIS images following treatment with either saline or 15mg/kg, 30mg/kg or 60mg/kg eliglustat via daily oral gavage. **G)** Relative fluorescence units (RFU) +/- StDev and weight (g) of BRCA1^co/co^; MMTV-Cre; p53^+/-^ tumors following eliglustat treatment. Statistical significance was determined by 1-sided Student T-test in comparison of eliglustat treatment relative to saline control. **H)** Tumor growth trajectories (% change from baseline (Day 1)) for BRCA1^co/co^; MMTV-Cre; p53^+/-^ tumor-bearing mice (N=3 per group) following treatment with either saline or 30mg/kg eliglustat via daily oral gavage. Kaplan-Meier survival curves are shown to the right. Statistical significance was determined by Log-rank (Mantel-Cox) tests and 2-sided p-value reported.

### Dose de-escalation studies and assessment of eliglustat-associated toxicity

The above investigations provide proof-of-concept for utility of eliglustat in the treatment of TNBC. To further assess translational potential of eliglustat, we performed a dose de-escalation study using an independent set of mice and additionally tested the anti-cancer efficacy of 15mg/kg and 30mg/kg eliglustat treatment, which are within range of approved clinical dosage in human patients adjusted for differences in species metabolic rates.^43,44^ Tumor growth was suppressed in a dose-dependent manner, reaching statistical significance (1-sided Welch Test p<0.05) at 30mg/kg (**Fig. 5F-G**). Prolonged treatment of TNBC-tumor bearing mice with 30m/kg eliglustat further demonstrated potent anti-cancer effects and markedly improved overall survival compared to mice treated with vehicle (saline) control (Log-rank (Mantel-Cox) test 2-sided p<0.001) (**Fig. 5H**).

No differences in body weight were found following intervention (**Fig. S10A**). Clinical chemistry tests following eliglustat treatment indicated dose-dependent elevations in blood ALT and AST levels. Blood calcium, chloride, potassium, sodium, lactate dehydrogenase, and glucose levels were impacted at 60mg/kg eliglustat treatment (**Fig. S10B**). Histopathological examination of kidney and liver from treated mice revealed no remarkable differences (all χ2 test for trend 2-sided *P-values* >0.1) compared to saline control (**Table S7**).

## DISCUSSION

The association between sphingolipids and mitochondria has, to date, primarily been considered in the context of ceramide-induced mitochondrial dysfunction. Here, we detail an additional, unrecognized role of sphingomyelin salvaging in mediating survival mitochondrial maintenance. Our findings integrate disparate previous concepts into a comprehensive model of extracellular sphingomyelin scavenging and intracellular salvaging directed towards glycosphingolipid biosynthesis that we demonstrate to specifically support the biogenesis of glycosphingolipid-enriched vesicles that provide an alternative route of mitochondrial turnover and clearance (**Fig. 6**). Importantly, we establish that this metabolic program is a targetable vulnerability in TNBC through repurposing of the selective glucosylceramide synthase inhibitor eliglustat, an FDA approved drug for long-term management of Gaucher Disease. Mechanistically, these studies demonstrate that eliglustat promotes accumulation of mitotoxic ceramides with subsequent mitochondrial damage and consequent cancer cell death. Moreover, we show that eliglustat suppresses tumor growth and drastically prolongs overall survival at clinical achievable doses in TNBC tumor-bearing mice.

**Figure 6.**
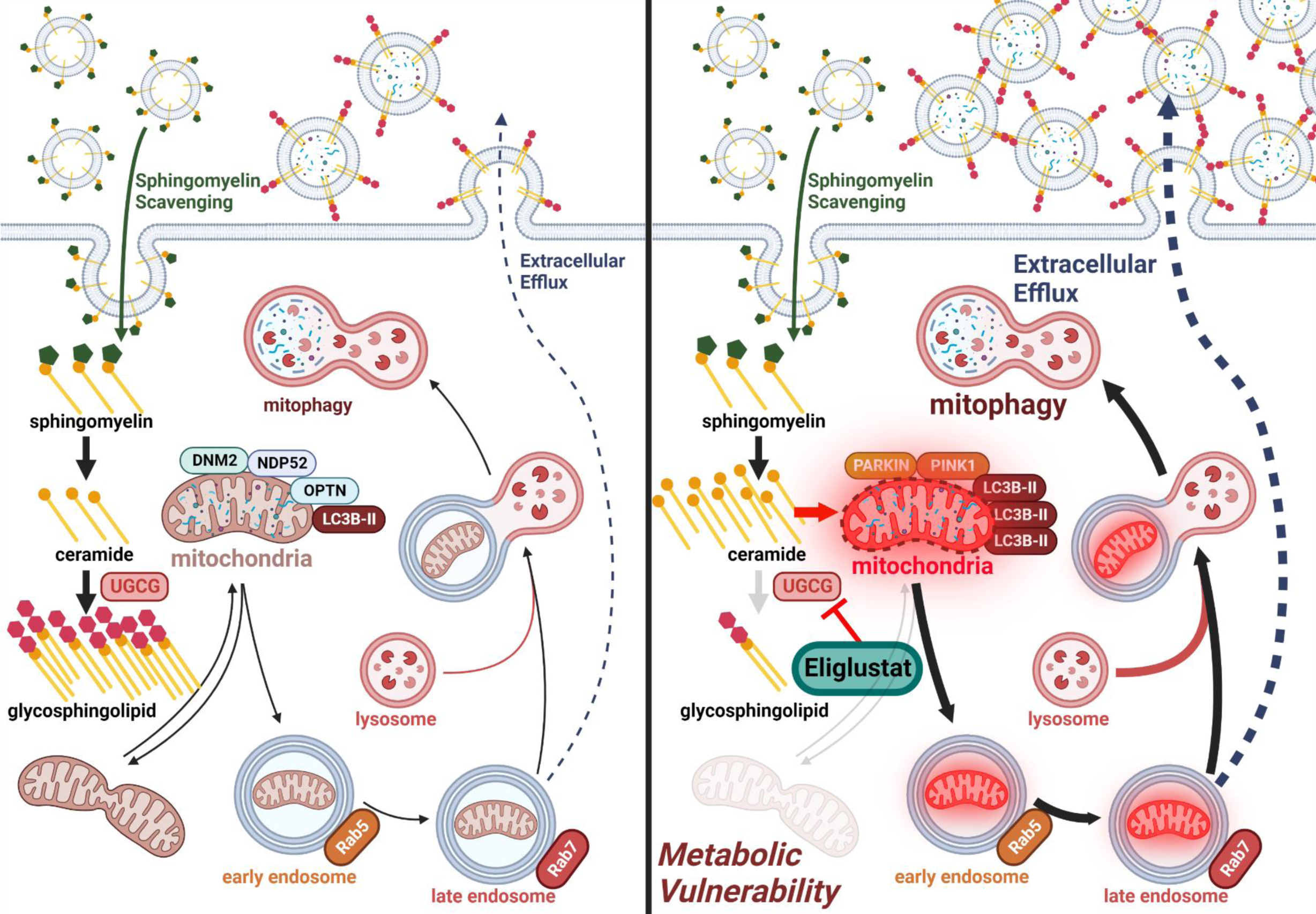
Proposed mechanism of sphingolipid oncometabolism. Essential components include (1) scavenging of sphingomyelin, (2) increased tumor catabolism of sphingomyelins to yield glycosphingolipids that support vesicle biogenesis that (3) intersects with ‘survival’ clearance of damaged mitochondria. *Metabolic Vulnerability:* Selective targeting of UGCG using small molecule inhibitors results in accumulation of mitotoxic ceramides in TNBC cells that promotes lethal mitophagy leading to direct anti-cancer effects as well as potentiation of the anti-tumor immune response.

Oncogenic stress-induced permeabilization of the outer mitochondrial membrane is known to induce intrinsic apoptosis.^45^ Mitochondria comprise approximately one-fifth of the cell cytoplasmic volume, and are structured into various networked and fragmented morphologies that are transitory and continuously altered through a balance of fission and fusion.^46^ Mitochondria associate with other organelles, and the mitochondrial-ER interface, known as the mitochondria-associated membrane (MAM), is a major hub of cell fate decisions. The lipid composition of MAMs is analogous to that of lipid raft domains, characterized by enrichment of distinct lipids, including sphingolipids and glycated derivatives. In addition, the ER network is elaborate, and the membranes of the ER cisternae engage in contact with other organellular membranes including those of the lysosomes, Golgi apparatus and endosomes as well as the plasma membrane.^47^ The lipid raft motif is conserved at these various membrane contact sites, facilitating organized compartmentalization of the intracellular milieu.^48^

Rather than relying on free diffusion, transfer of lipids and other metabolites to, within, and out of mitochondria is increasingly understood to entail coordinated trafficking between mitochondrial and other organellar membranes via mechanisms that include direct contact as well as protein- and vesicle-mediated transport.^49^ Recent evidence indicates that there is extensive biochemical crosstalk between mitochondria and other organelles, and mitochondrial-derived vesicles (MDVs) have emerged as an important transporter of molecular cargoes from mitochondria. Distinct fates for MDVs including targeting to either to the late endosome/multivesicular body or to peroxisomes have been identified, and MDV biogenesis has been found to involve Vps35, Parkin, and PINK1, proteins also known to play a critical role in Parkinson’s disease pathogenesis.^10^ It is becoming increasingly recognized that the specific distribution and local concentration of lipid components at these interfaces is instrumental in defining their physiological function and that alterations in recruitment and turnover of requisite lipid species to these sites is an underlying feature in numerous disease pathologies. Here we provide evidence of a metabolic network of sphingolipid scavenging and catabolism that intersects with vesicle biogenesis and trafficking to enable survival clearance in the context of triple-negative breast cancers.

The development of additional targeted treatment modalities that elicit potent anti-cancer effects while yielding favorable toxicity profiles is clinically desirable. Our investigations demonstrate that targeting of the sphingolipid metabolic network described here via the selective glucosylceramide synthase inhibitor eliglustat elicits anti-cancer effects by promoting accumulation of mitotoxic ceramides, induction of lethal mitophagy and subsequent cancer cell death both *in vitro* and *in vivo*. Dose de-escalation studies demonstrated that eliglustat anti-cancer effects are achievable at clinically relevant lower doses with favorable toxicity profiles. These findings, coupled with the fact that eliglustat was developed for long-term treatment of Gaucher Disease and has been shown to be well tolerated in these patients, provides strong rationale for the application of eliglustat as an anti-cancer agent for the treatment of TNBC.

In summary, we define a mechanism of aberrant sphingolipid metabolism in TNBC that intersects with mitochondrial dynamics and vesicle biogenesis. We further demonstrate that this activity presents a metabolic vulnerability that can be directly targeted by repurposing eliglustat, an existing selective inhibitor of glucosylceramide synthase. Additional studies are warranted to further develop the clinical utility of these findings in the context of TNBC.

## Acknowledgments

This work was Supported by the generous philanthropic contributions to The University of Texas MD Anderson Cancer Center Moon Shots Program™, the Little Green Book Foundation, and the Brown Foundation. J.F.F. was supported by an NCI 1R37CA269611-01A1.

## Author contributions

Conceptualization: JV, JFF

Methodology: JV, HK, EI, RW, EM, JBD, APH, LR, YC, FCH, JFF

Investigation: JV, YC, MH, MZ, JFF

Visualization: RAL, JFF

Data Analysis: EI, JFF

Resources: ABG, BA, SH, JFF

Funding acquisition: SH, JFF

Project administration: JBD, SH, JFF

Supervision: JFF

Writing – original draft: JFF

Writing – review & editing: JV, MJH, MZ, HK, HK, EI, RW, EM, JBD, ABG, APH, TCT, PB, LR, YC, FCH, SP, BA, SH

## Competing Interests

Authors declare no conflicts-of-interest.

## Notes

**Conflict-of-interest:** The authors declare no potential conflicts of interest.

### Competing Interest Statement

The authors have declared no competing interest.

