## Supplementary material for "Vesicle-mediated mitochondrial clearance underlies an actionable metabolic vulnerability in triple-negative breast cancer": Supplemental Materials.docx

### **Supplemental Table S1-7**

***See excel files***

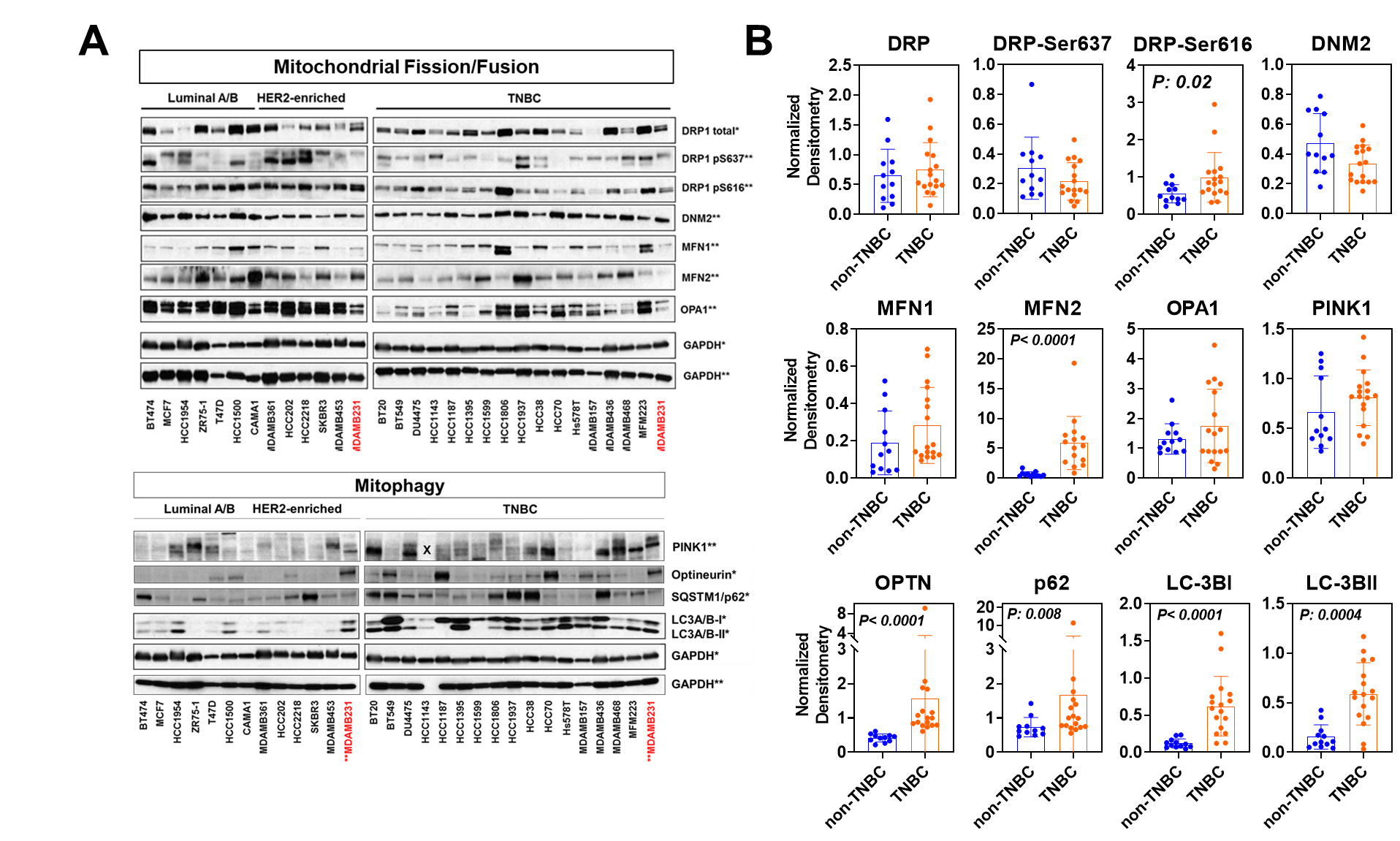

**Supplemental Figure S1. Expression patterns of autophagy and mitophagy-related proteins across breast cancer cell lines. A)** Immunoblots illustrating expression levels of mitochondrial fission and fusion-related proteins as well as mitophagy related proteins among 28 breast cancer cell lines. MDAMB231 was used as a reference control among the different gels. PINK1 was not able to be assessed for HCC1143 in gel #2 (annotated as “X”). **B)** Normalized abundances (densitometry). Densities for a given protein were first normalized against respective GAPDH (loading control). To account for differences in signal exposure, GAPDH-normalized bands were subsequently standardized against the MDAMB231 reference bands. Statistical significance was determined by Wilcoxon rank sum test and 2-sided p-values reported.

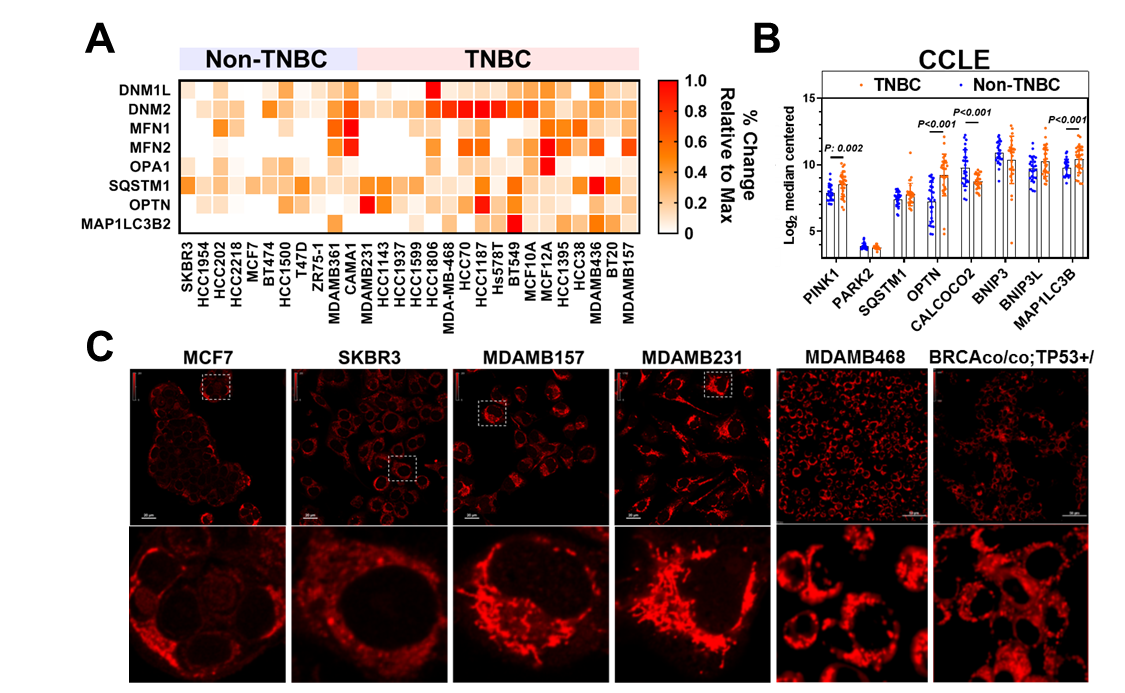

**Supplemental Figure S2. Gene expression of mitophagy related genes among TNBC and non-TNBC cell lines from the Cancer Cell Line Encyclopedia (CCLE) transcriptomic database.** Statistical significance was determined by Wilcoxon rank sum test and 2-sided p-values reported.

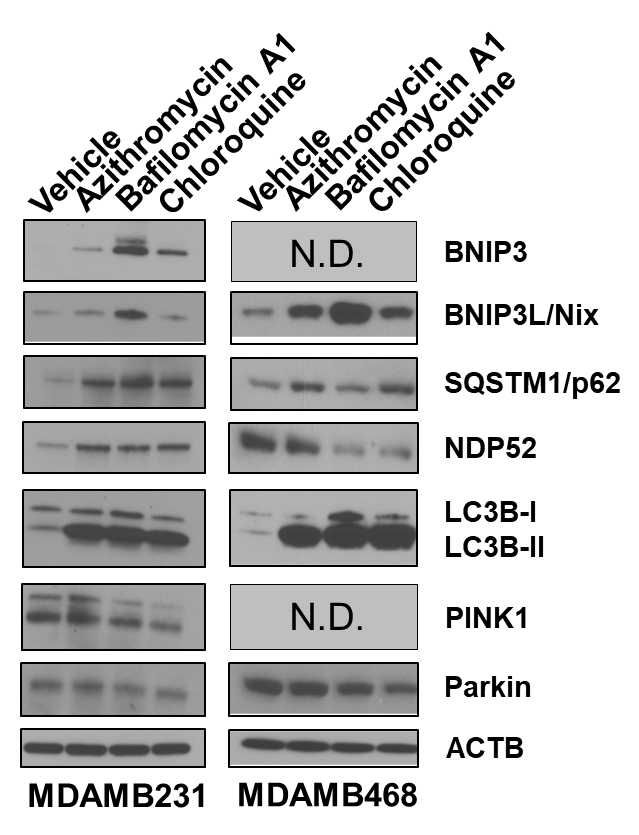

**Supplemental Figure S3. Immunoblots for mitophagy-related proteins in MDAMB231 and MDMB468 TNBC cells following 24-hour treatment with vehicle (DMSO) control, 10uM azithromycin, 1nM bafilomycin, or 10uM Chloroquine.** N.D.- not detected

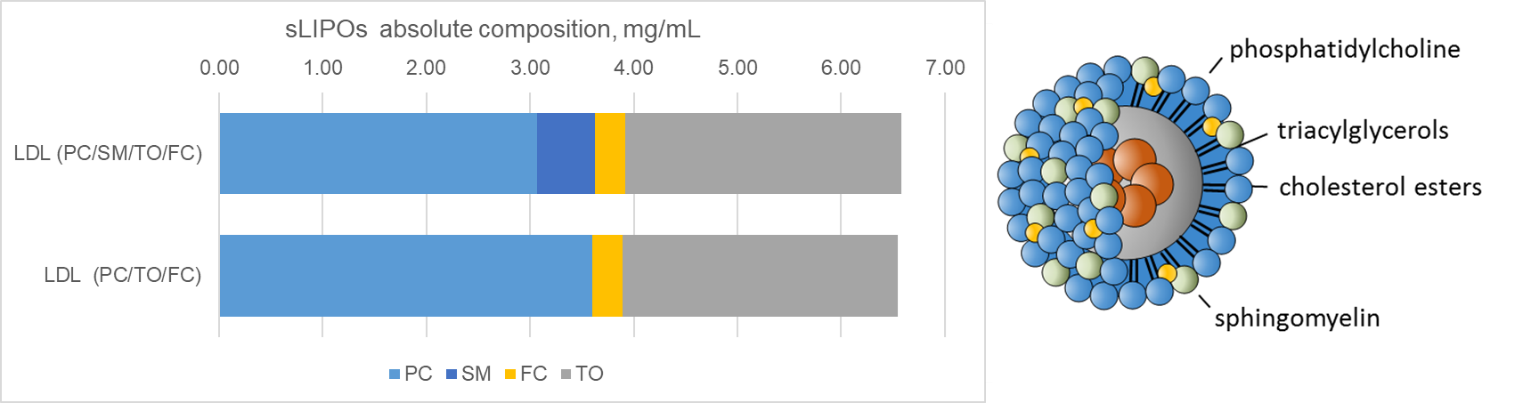

**Supplementary Figure S4. Schematic depicting composition of synthetic self-assembled lipid particles (SSALPs).** PC- phosphatidylcholine; SM- sphingomyelin; FC- free cholesterol; TO-trioleate

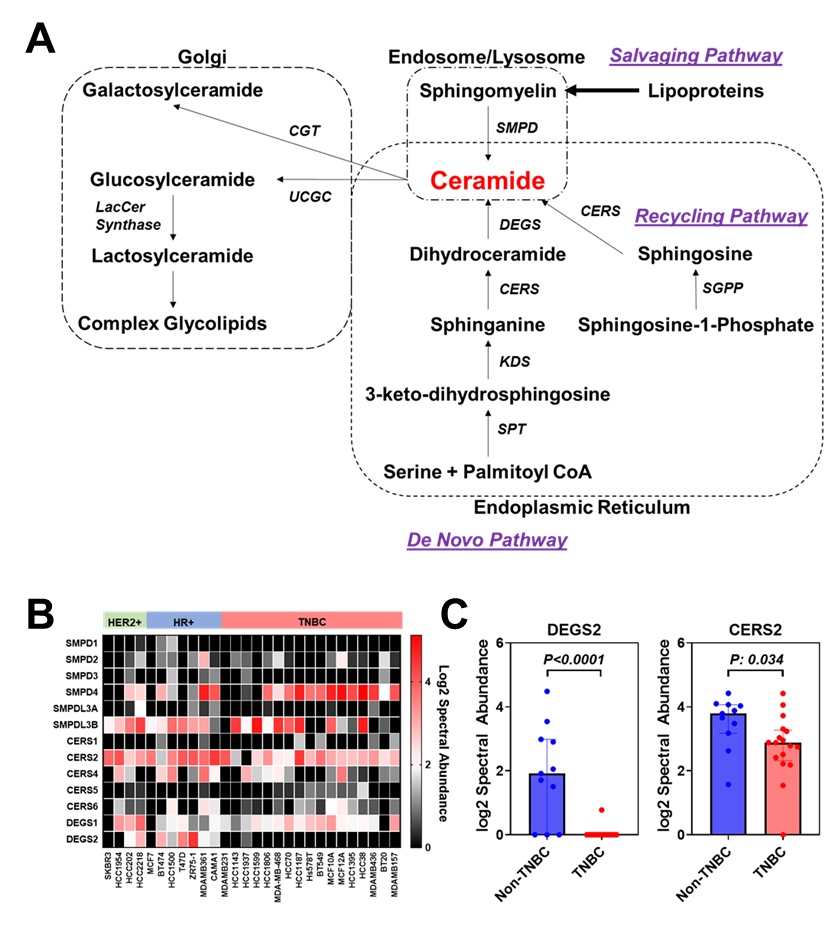

**Supplemental Figure S5. Protein levels of enzymes involved in ceramide metabolism in TNBC and non-TNBC cell lines. A)** Schematic of ceramide biosynthetic pathways. **B)** Heatmap illustrating whole cell lysate protein expression (log2 spectral abundance) of sphingolipid-metabolizing enzymes among 28 breast cancer cell lines. **C)** Spectral abundance of DEGS2 and CERS2 in non-TNBC and TNBC cell lines. Statistical significance was determined by 2-sided Wilcoxon rank sum test.

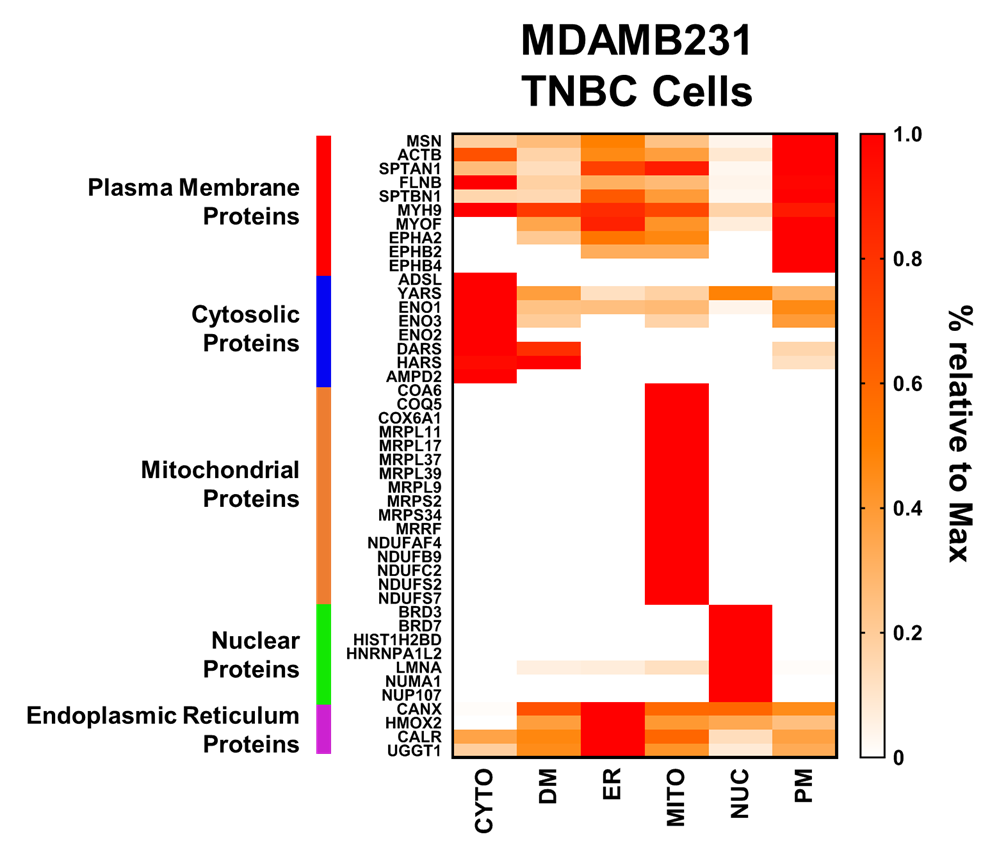

**Supplementary Figure S6. Heatmap depicting organelle-enriched markers in subcellular fractions of MDAMB231 TNBC cells.** Values represent % relative to the maximum observed abundance for the given protein across the different subcellular compartments. Abbreviations. **CYTO-**cytoplasm; **DM-** Dense microsomes; **ER-** endoplasmic reticulum; **MITO-** mitochondria; **NUC-** nuclear compartment; **PM-** plasma membrane.

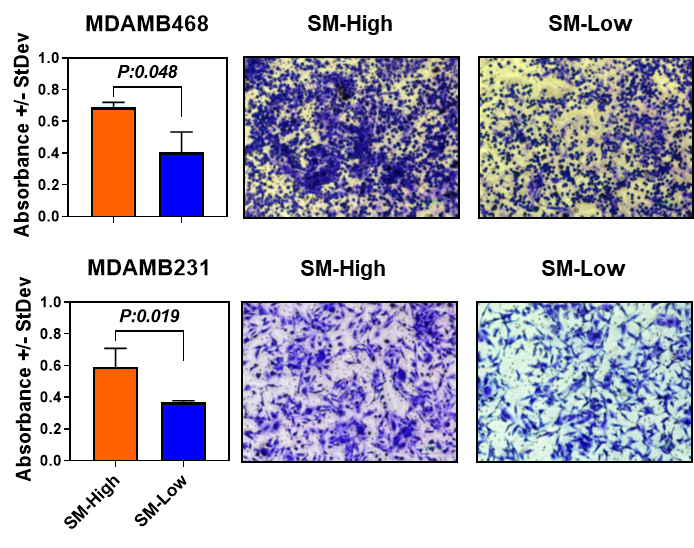

**Supplementary Figure S7. Extracellular sphingomyelin deprivation attenuates TNBC cell invasiveness.** MDAMB231 and MDAMB468 TNBC cells were treated with sphingomyelin (SM)-high or SM-low self-assembled lipid particles and TNBC cell invasive properties assessed using cell invasion assays. Cells in the bottom chamber were stained with crystal violet and representative images and quantitative readouts (absorbance) provided. Statistical significance was determined by 1-sided Student T-test.

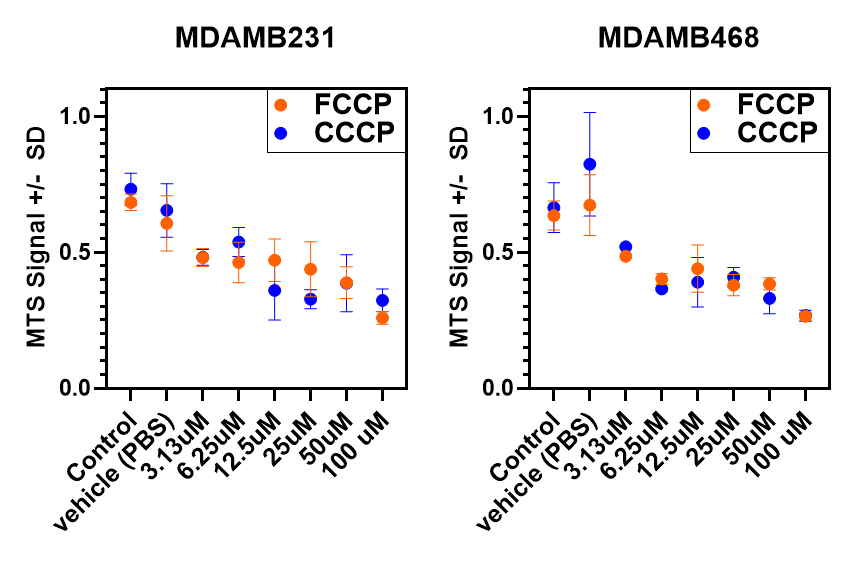

**Supplemental Figure S8. Viability curves for MDAMB231 and MDAMB468 TNBC cells following treatment with FCCP or CCCP.**

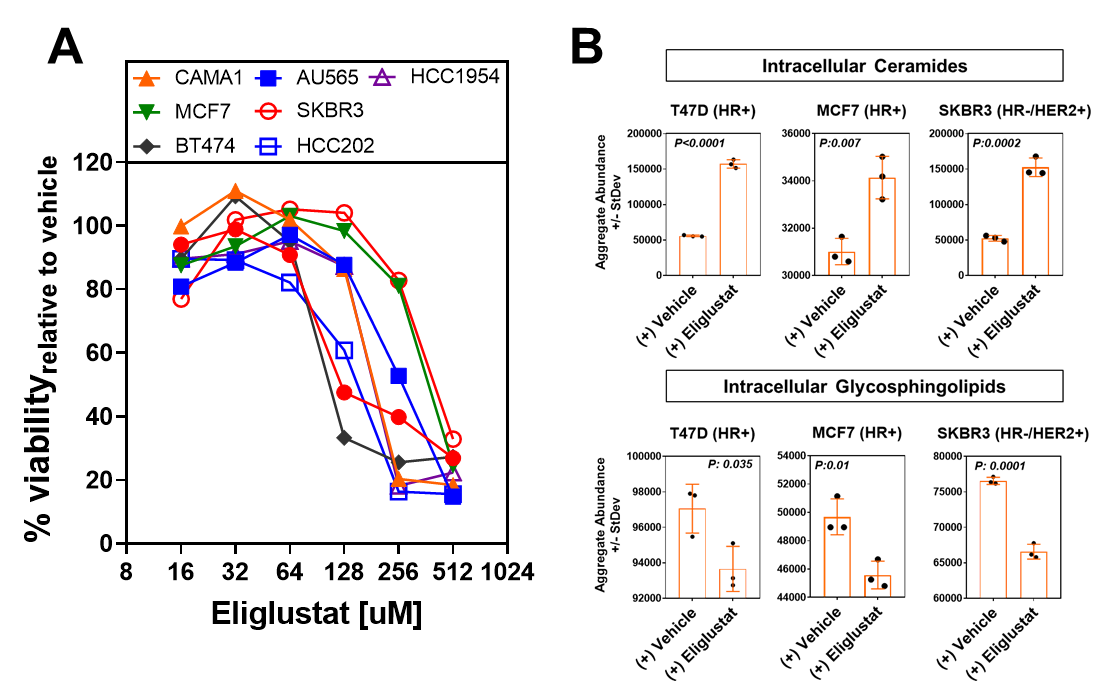

**Supplemental Figure S9. Anti-cancer effects of eliglustat in non-TNBC breast cancer cell lines. A)** Viability curves for a panel of 7 non-TNBC cell lines following treatment with eliglustat. Viability was determined by MTS assay. **B)** whole cell lysate levels of ceramides (left panel) and glycosphingolipids (right panel) in human non-TNBC cells following 6-hour eliglustat treatment. Values represent aggregate area abundances of ceramides or glycosphingolipids detected by mass spectrometry. Statistical significance was determined by 1-sided Student T-tests.

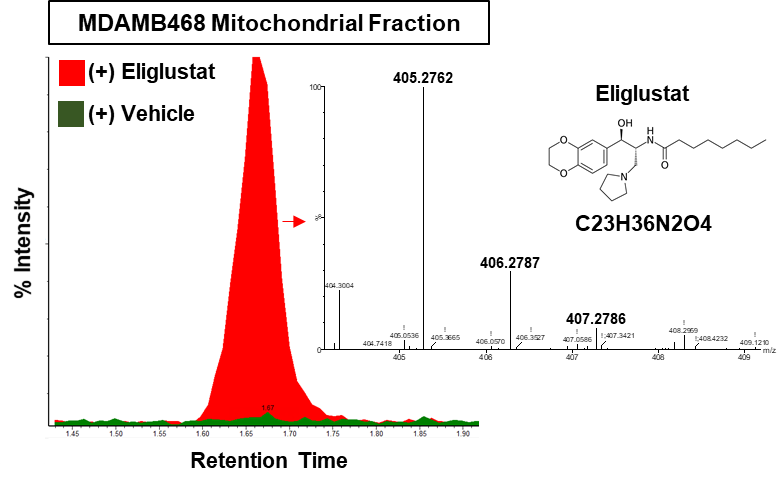

**Supplemental Figure S10. Extracted ion chromatogram for eliglustat in mitochondria isolated from MDAMB468 TNBC cells following treatment with eliglustat or vehicle control.** The [M+H]+ ion (m/z 405.2762) for eliglustat is shown.

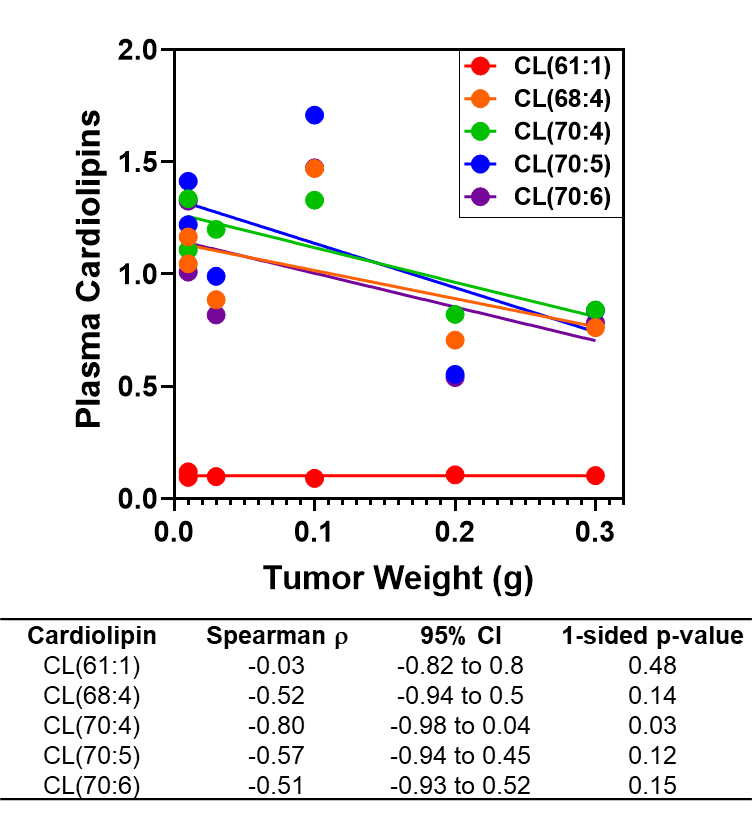

**Supplemental Figure S11. Association between plasma cardiolipins and tumor weight in BRCA1co/co; MMTV-Cre; p53+/- TNBC tumor-bearing mice following eliglustat treatment.**

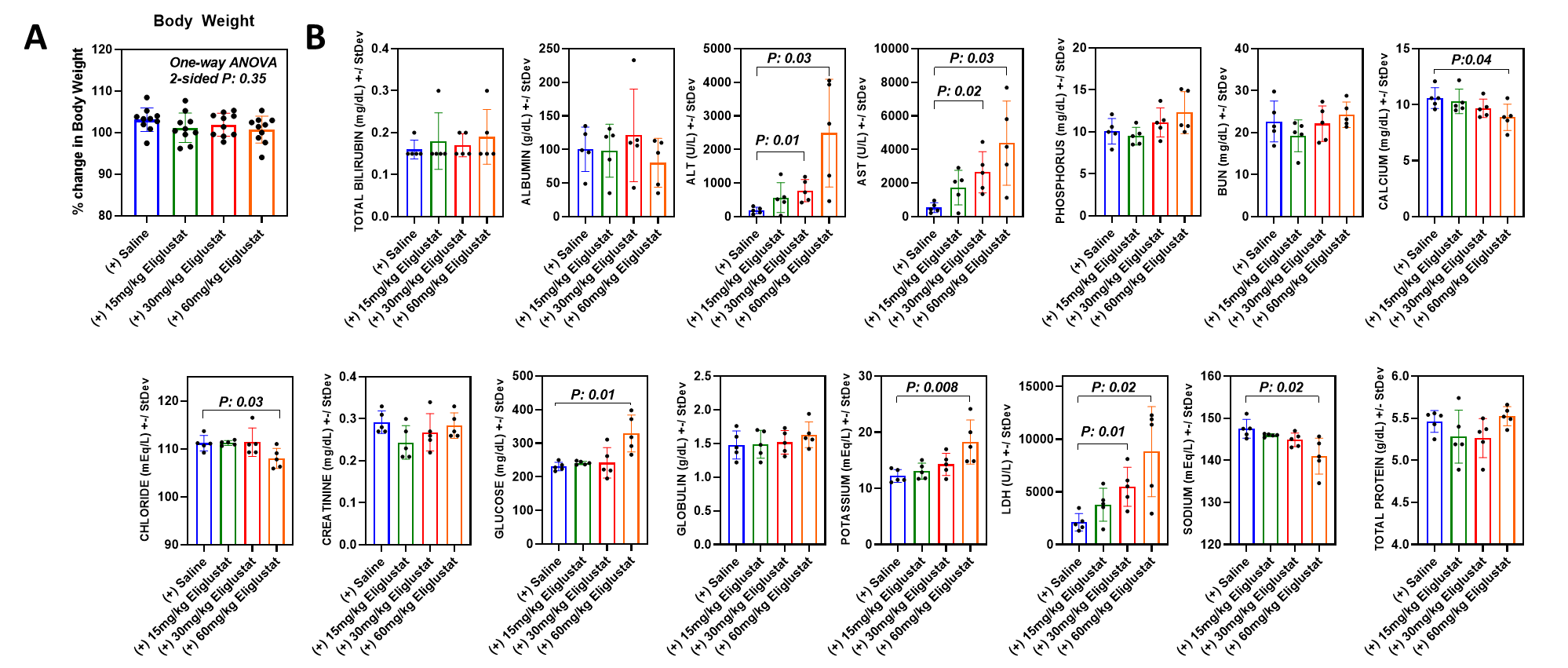
**Supplemental Figure S12. Body weight, blood chemistry, and kidney and liver histopathological results following eliglustat treatment in BRCA1co/co; MMTV-Cre; p53+/- TNBC tumor-bearing mice. A)** % change in body weight following intervention. Statistical significance was determined by One-Way ANOVA and a 2-sided P-value is reported. **B)** Blood chemistry results following intervention. Statistical significance was determined by 2-sided Welch T-test in comparison to saline control. N=5 mice per treatment condition.

### **References**
